# Molecular phenotyping of single pancreatic islet leader beta cells by “Flash-Seq”

**DOI:** 10.1101/2022.08.26.505442

**Authors:** Pauline Chabosseau, Fiona Yong, Luis F. Delgadillo-Silva, Eun Young Lee, Shiying Li, Nidhi Gandhi, Jules Wastin, Livia Lopez Noriega, Isabelle Leclerc, Yusuf Ali, Jing W. Hughes, Robert Sladek, Aida Martinez-Sanchez, Guy A. Rutter

**Affiliations:** Centre de Recherche du CHUM, Faculté de Médicine, Université de Montréal, Montréal, QC, Canada; Section of Cell Biology and Functional Genomics, Division of Diabetes, Endocrinology and Metabolism, Department of Metabolism, Digestion and Reproduction, Faculty of Medicine, Imperial College London, London W12 0NN, United Kingdom; Lee Kong Chian Imperial Medical School, Nanyang Technological University, Singapore; Department of Medicine, Washington University School of Medicine, Saint Louis, MO, United States; Division of Endocrinology and Metabolism, Department of Internal Medicine, Seoul St. Mary’s Hospital, The Catholic University of Korea, Seoul, South Korea; Departments of Medicine and Human Genetics, McGill University and Genome Quebec Innovation Centre, Montreal, QC, Canada

**Author notes:** Please address correspondence to Professor Guy Rutter.

**Keywords:** diabetes, beta cell, calcium signalling connectivity, insulin secretion, RNA-Seq

## Abstract

**Aims:** Spatially-organised increases in cytosolic Ca^2+^ within pancreatic beta cells in the pancreatic islet underlie the stimulation of insulin secretion by high glucose. Recent data have revealed the existence of subpopulations of beta cells including “leaders” which initiate Ca^2+^ waves. Whether leader cells possess unique molecular features, or localisation, is unknown.

**Main methods:** High speed confocal Ca^2+^ imaging was used to identify leader cells and connectivity analysis, running under MATLAB and Python, to identify highly connected “hub” cells. To explore transcriptomic differences between beta cell sub-groups, individual leaders or followers were labelled by photo-activation of the cryptic fluorescent protein PA-mCherry and subjected to single cell RNA sequencing (“Flash-Seq”).

**Key findings:** Distinct Ca^2+^ wave types were identified in individual islets, with leader cells present in 73 % (28 of 38 islets imaged). Scale-free, power law-adherent behaviour was also observed in 29% of islets, though “hub” cells in these islets did not overlap with leaders. Transcripts differentially expressed (295; padj<0.05) between leader and follower cells included genes involved in cilium biogenesis and transcriptional regulation. Functionally validating these findings, cilia number and length tended to be lower in leader *vs* follower cells. Leader cells were also located significantly closer to delta cells in Euclidian space than were follower cells.

**Significance:** The existence of both a discrete transcriptome and unique localisation implies a role for these features in defining the specialized function of leaders. Specifically, these data raise the possibility of altered signalling from delta cells towards somatostatin receptors present on leader cell cilia.

## INTRODUCTION

Adequate insulin secretion is necessary to regulate blood glucose levels efficiently in mammals, and failure to maintain sufficient insulin secretion in the context of insulin resistance is at the core of most forms of type 2 diabetes (T2D) [1]. Insulin-secreting beta cells reside within the islets of Langerhans, endocrine micro-organs scattered across the pancreas, and make up ~60 % in human and 85% of total islet cells in rodents [2].

There is considerable evidence to suggest that beta cells are not a homogeneous population. Thus, variations between individual cells exist in insulin secreting capabilities and sensitivity to glucose [3–7] as well as at the level of the transcriptome [8–15]. Moreover, we [16,17] and others [18,19] as reviewed in Korošak et al [20], have shown that specific beta cell subpopulations play discrete and definable roles within the intact islet, coordinating the overall response to stimulation with glucose or other secretagogues [21,22]. Importantly, artificial lowering of heterogeneity across the islet, achieved by forced overexpression of *MafA* and *Pdx1* compromises insulin secretion [23].

Using high-speed calcium imaging we have previously identified a subgroup of highly connected cells named “hubs” [16]. This population corresponds to a small proportion (~10%) of cells which, when examined using an optogenetics approach, were shown to control calcium responses across the islet [16,17]. Hub cells were shown to display relatively higher levels of glucokinase, and lower insulin, immunoreactivity, than follower cells [16], allowing the imputation of transcriptomic differences between the two groups through analysis of previously published single cell RNA data sets [17]. Other sub-categories include “wave originator” cells, from which calcium waves propagate in response to glucose during the first acute insulin secretion phase (also called “first responders” in response to a change from low to high glucose). During the sustained, plateau phase of the response, “leader” cells represent those from which calcium waves emanate [16,17,24]. Whether or not leader cells, first responders and hub cells are fully distinct, or represent partly overlapping subpopulations, is presently unclear [24].

An important advance in our understanding of the roles of these cells would be made by ascribing molecular signatures (e.g. transcriptomic, metabolomic or proteomic) to each group after their identification – based on function – and subsequent isolation. With this goal in mind, we first demonstrate that, in mouse islets displaying Ca^2+^ oscillations at constant glucose (11mM), repetitive Ca^2+^ waves usually emanate from a main stable leader cell. Next, by performing connectivity analysis we reveal that hubs and leaders are distinct subpopulations. Thirdly, we used targeted photopainting of individual cells, followed by RNA sequencing (“Flash-Seq”), to demonstrate that leader cells possess a discrete transcriptome versus “follower” cells. Differentially expressed genes included transcriptional regulators and several encoded proteins involved in cell cilium biogenesis and assembly.

## RESULTS

### Repetitive Ca^2+^ waves usually emanate from a stable leader cell within the islet

In order to identify and subsequently isolate individual beta cells with distinct roles in the generation and transmission of Ca^2+^ waves, we used islets bearing the recombinant Ca^2+^ probe GCaMP6f and photo-activatable PA-mCherry (Methods). Ca^2+^ imaging was performed at a stimulatory but submaximal glucose concentration (11 mM), with image acquisition at 5 Hz in a single plane (Figure 1A,B – Supplementary movie 1). Data were analysed from 43 islets imaged in five independent experiments involving a total of nine separate islet preparations. Of all the islets interrogated, three were unresponsive to glucose stimulation, two displayed uncoordinated responses (where cells showed individual calcium oscillations without propagation to neighbours), and 38 showed coordinated Ca^2+^ oscillations where signal propagation mobilised >60% of cells through the islet (Figure 1C). Spatially, the observed oscillations varied in terms of propagation rate and duration. On average, islets oscillated at a rate of 0.86±0.23 oscillations min^-1^, with 27 islets showing sustained oscillations (whose duration was > 5 s from the instant 50% of total cells showed an increased fluorescence signal – higher than 50% of baseline – to that at which the signal in 50 % of cells had declined to baseline), and 11 islets quick oscillations (duration <4.9 s; Figure 1D,E, Table 1, Supplementary Movie 2-3).

**Figure 1.**
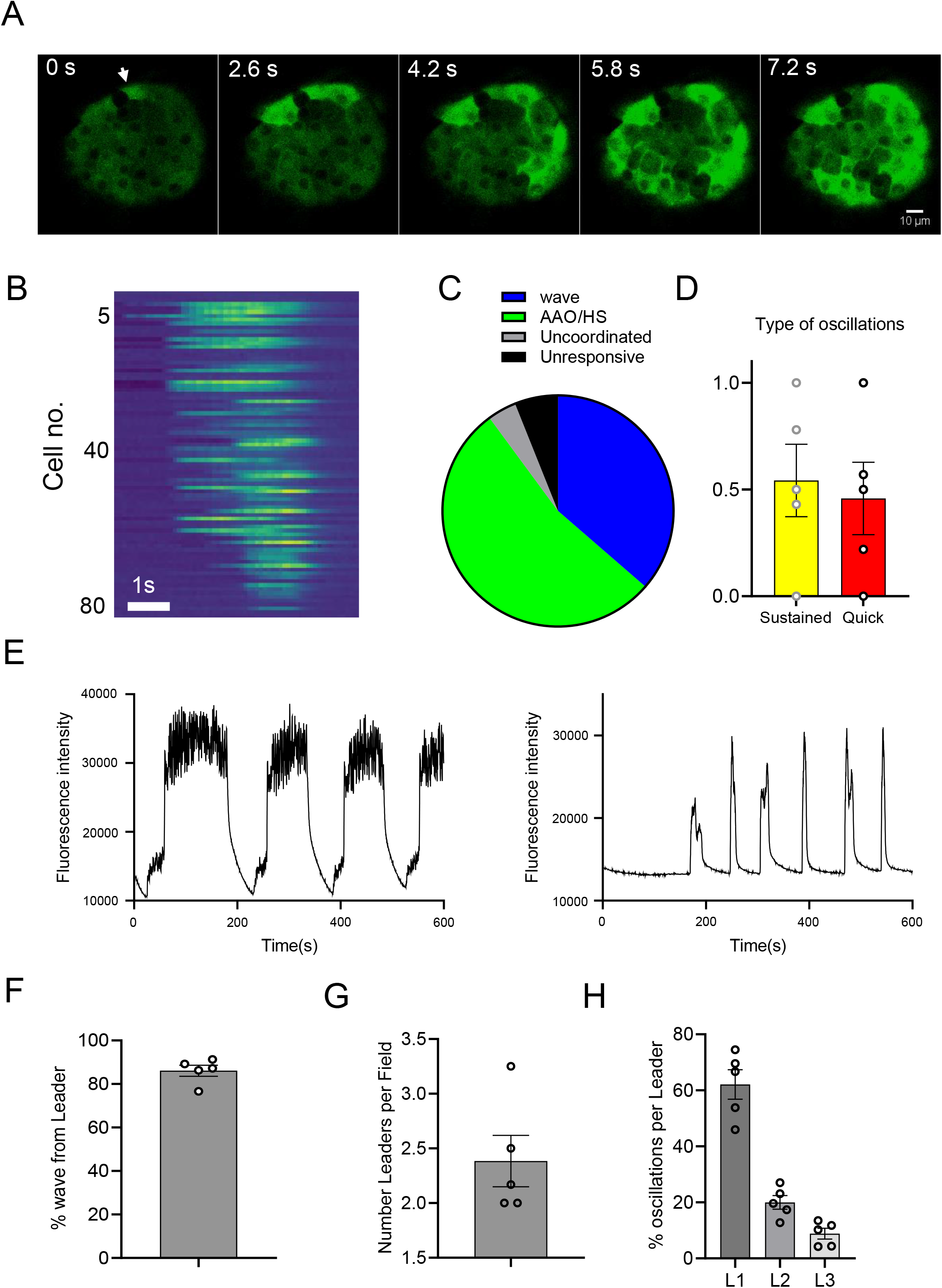
Islet Ca^2+^ wave properties. **A.** Timelapse of a calcium wave propagation through the islet during an oscillation in response to glucose at 11 mM. The leader cell (arrow) is the first cell to show an increase in fluorescence signal, which then propagates to surrounding neighbour cells until all the cells show an increased fluorescence signal corresponding to calcium entry, step leading to insulin secretion. **B**. Raster plot corresponding to the acquisition shown in A and Supplementary Movie 1. Cell “5” corresponds to the arrowed cell in A. **C.** Categorisation of islets depending on the signal propagation behaviour during calcium waves. 6% were unresponsive, 4% displayed uncoordinated response ie. no wave, 36% slow propagation wave (>1 s) and 53 % quick propagation wave (All-at-once/High-speed AAO/HS – <1s) n= 43 islets from 5 experiments **D.** Categorisation of islets depending on the lasting time of oscillations. After wave propagations, the time of activation of each oscillation was measured and averaged for each islet showing a waving behaviour (38 islets – 5 experiments). Islets for which activation were >5s were classified in the category of sustained oscillations, and <5s were classified as quick oscillations **E.** Representative trace for islets with sustained oscillation (left panel) and islets with quick oscillations (right panel). **F.** Leader cells were defined as being the first cell to show a signal increase during an oscillation. In the great majority of cases (from 80 to 95% – 38 islets from 5 experiments) leader cells were wave originator, ie signal would propagate from the leader cells to the direct neighbouring cells. **G.** For each islet with a waving behaviour, number of leader cells was determined during time the time of acquisition (10 minutes) and were in average from 2 to 4 cells (28 islets – 5 experiments) **H**. During acquisition time, islets had in average 1 oscillation per minute. Repartition of waves between leader cells were quantified. In average, one main leader cell was responsible of 61±6% of the waves, while a secondary one 29±9.6%, and third ~10%.

**Table 1.** Islets classification into subgroups based on Ca^2+^ signal propagation rate behaviour and oscillation behaviour.

|  | Wave | AAO | Total number of islets | Power Law compliance |
| --- | --- | --- | --- | --- |
| Quick | 7 | 4 | 11 | Y |
| Sustained | 9 | 18 | 27 | N |
| Total number of islets | 16 | 22 | 38 |  |

Two types of Ca^2+^ signal propagation types were identified, and islets were placed into one of two categories accordingly. The first included islets where the propagation time of a Ca^2+^ signal from the wave originator cell to neighbouring cells – to the point at which >70% cells displayed an increase in Ca^2+^ signal to 50% above baseline (Figure 1A,B, from first image to last image, Supplementary Movie 4) – was >1s. In the second, where the propagation time was ≤1s, islets were categorised as displaying an “all-at-once”/high-speed wave (AAO/HS) behaviour (Supplementary Movie 5).

Leader cells were defined as those exhibiting the first detectable increase in Ca^2+^ signal (50% above baseline) during Ca^2+^ waves, and were detected in 73±4% of the islets imaged (28 islets out of 38 in total, from five independent experiments). Depending on the oscillation rate of each islet (i.e., the number of oscillations per minute, observed during the 10 min. acquisition), 1 - 4 leader cells were observed, with an average of 2.4±0.23 leader cells per plane of view (Figure 1F; n= 28 islets from five independent experiments). The vast majority of leader cells (86.1±2.5%) acted as wave originators (Figure 1G). Strikingly, the distribution of waves between wave originator cells was not even. Thus, the majority of the Ca^2+^ waves (61±6%) emanated from a “principal” leader cell, while a “secondary” leader cell was responsible for 29±9.6% of Ca^2+^ waves. The remaining ~10% of waves started from a third leader cell (Figure 1H).

### Connectivity analysis reveals hubs and leaders are distinct subpopulations

Islets were next subdivided into four subgroups based on Ca^2+^ signal propagation rate and oscillation behaviour (Figure 1, Table 1).

We next explored beta cell-beta cell connectivity [16,17,22,24]. Figure 2A shows the Ca^2+^ fluorescence traces and topographic representations of the connected beta cells for each subgroup based on the strength of the coactivity between any two cells. Cells are represented by differently coloured nodes depending on their coactivity: black indicates cells that coactivate with ≥80% of the remaining beta cells, while grey and white nodes represent beta cells that coactivate with ≥60% and ≥40% respectively with the rest of the beta cell population. Nodes circled with a solid black line indicate leader cells, as defined above (see Supplementary Movie 2-5).

**Figure 2.**
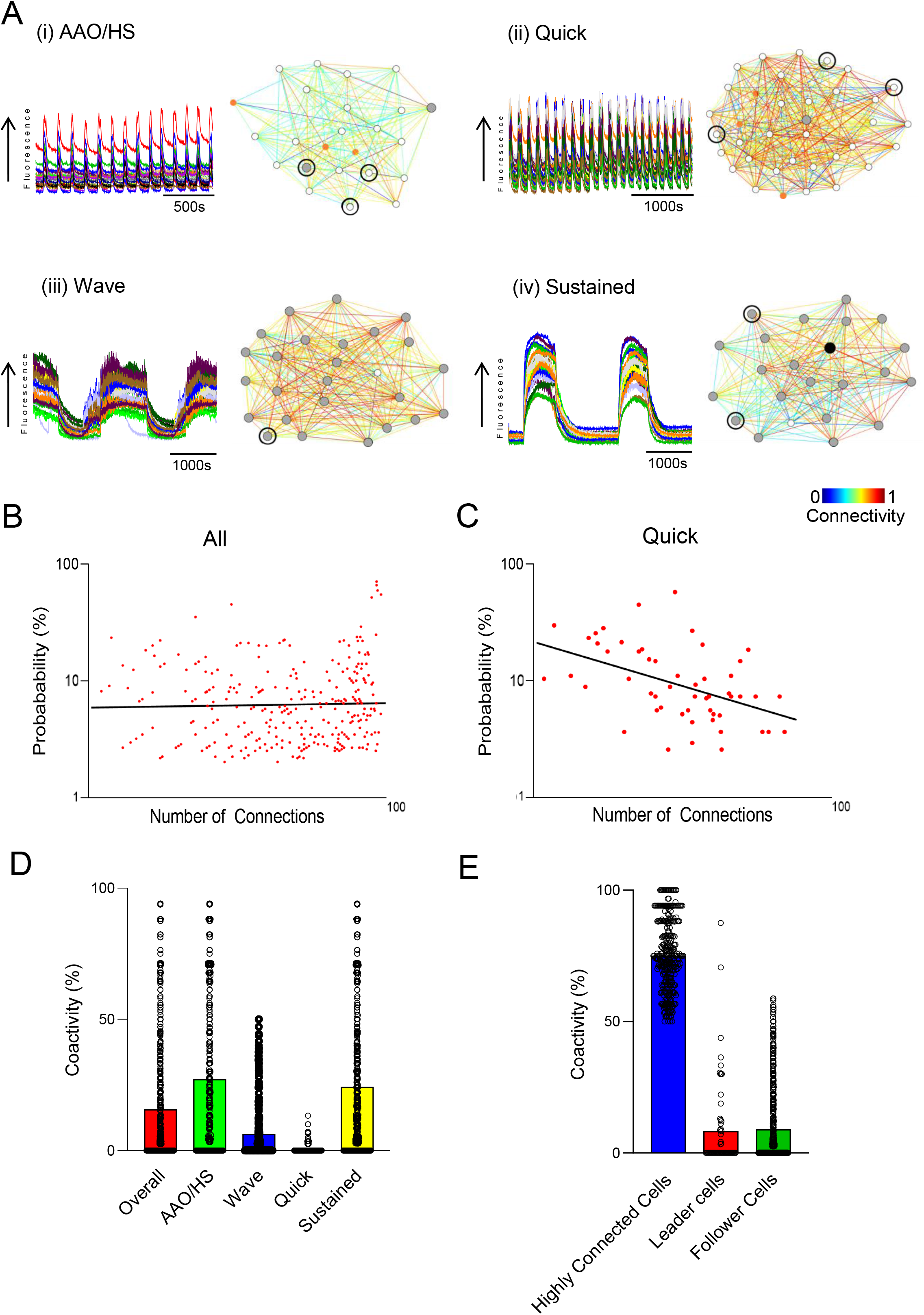
Intra-islet connectivity. **A.** Fluorescence intensity readouts for all beta cells identified during a typical Ca^2+^ wave (superimposed in different colours) in a single representative islet for each subgroup. (i. AAO/HS, ii. Quick, iii. Wave, iv. Sustained) Representative cartesian maps of islets with colour coded lines connecting cells according to the strength of coactivation (colour coded R values from 0 to 1, blue to red respectively). **B.** Log-log graph of the beta cell-beta cell connectivity distribution (n=38). 37.2% of beta cells coactivated with at least 50% of the rest of the beta cells, identifying more than a third of all beta cells to be highly connected cells. **C.** Log-log graph of the beta cell-beta cell connectivity distribution in the quick subgroup. 1.86% of beta cells coactivated with at least 50% of the rest of the beta cells, identifying hub cells that display an obedience to a power-law distribution whereby few beta cells host 50% to 100% of the connections to the rest of the beta cell population. **D.** Average beta cell-beta cell coactivity overall and in each subgroup. A one-way ANOVA showed that each subgroup was significantly different in terms of their respective intra-islet beta cell coactivity (p≤0.01). However, there were no significant differences between the AAO and Sustained subgroups (p=0.27). **E.** Percentage coactivity of beta cells. Identified highly connected cells displayed coordinated Ca2+ responses to an average of 75.3% of all beta cells whereas identified leader and follower cells had significantly fewer coordinated Ca^2+^ responses at averages of only 8.35% and 9.70% respectively (p<.001). There were no significant differences in the average coactivity between the leaders and follower cells (p=0.72).

Unexpectedly, pooled data taken over 38 islets together did not reveal a scale-free network topography (Figure 2B), in contrast to previous studies [16,17,22,24], and we were unable to identify a small proportion (5% to 10%) of highly connected “hub” cells in these islets. Nevertheless, connectivity analysis revealed that more than a third (37.2±6.34%) of beta cells were highly connected, as assessed by counting the number of beta cells displaying coordinated Ca^2+^ responses with at least 50% of all beta cells. This finding also applied when subgroup analyses were performed across different wave types (Supplementary Figure 1), with one exception. Thus, the 11 islets that exhibited quick waves demonstrated scale-free network topography in which ~2% of the beta hub cells hosted >50% of the connections (Figure 2C; R^2^=0.17).

Beta cells displayed an average of 15.8% coactivity with all beta cells within all islets examined (Figure 2D) with the subgroups displaying a wide range of average coactivities: The “Quick” subgroup displayed 0.48% coactivity whereas this value was 27.3% in the “AAO” subgroup (Figure 2D; overall: 15.8%; AAO: 27.3%; Wave: 6.36%; Quick: 0.48%; Sustained: 24.3%). These differences were confirmed by one-way ANOVA (Figure 2D; p≤0.01), with the exception of the AAO versus Sustained subgroups (p=0.27).

Overall, highly connected beta cells (including “hubs” present in the Quick subgroup) displayed an average of 75.3% coactivity with all beta cells (Figure 2E). In contrast, identified leaders and follower cells had significantly lower coactivity, being linked to an average of only 8.35% and 9.70% of all beta cells respectively (Figure 2E; p<.001). There were no significant differences in the average coactivity between the leader and follower cells (Figure 2E, p=0.72).

Similar to the global analysis across all islets, highly connected cells displayed significantly higher coactivity on average in comparison to leaders (Supplementary Figure 2; p<.001; AAO: 77.7% vs 15.9%; Wave: 69.4% vs 4.13%; Quick: 54.1% vs 0.15%; Sustained: 75.6% vs 13.8%). There was also significantly more coactivation on average in comparison to follower cells (Supplementary Figure 2; p<.001; AAO: 77.7% vs 14.9%; Wave: 69.4% vs 6.57%; Quick: 54.1% vs 0.51%; Sustained: 75.6% vs 15.5%). There were no significant differences in the average coactivity between the leader and follower cells across all subgroups (Supplementary Figure 2; AAO: p=0.55; Wave: p=0.20; Quick: p=0.50; Sustained: p=0.75).

We next explored the possibility that different subgroups of cells may be more or less likely to trigger a subsequent Ca^2+^ wave. Multivariate vector autoregression (MVAR) analysis found causal relationships between beta cells in all glucose-responsive islets. However, MVAR leaders were found in only 31.6% of the 38 responsive islets. The Wave subgroup made up half of these islets (Figure 3A). MVAR leaders were also found in each subgroup (Figure 3B).

**Figure 3.**
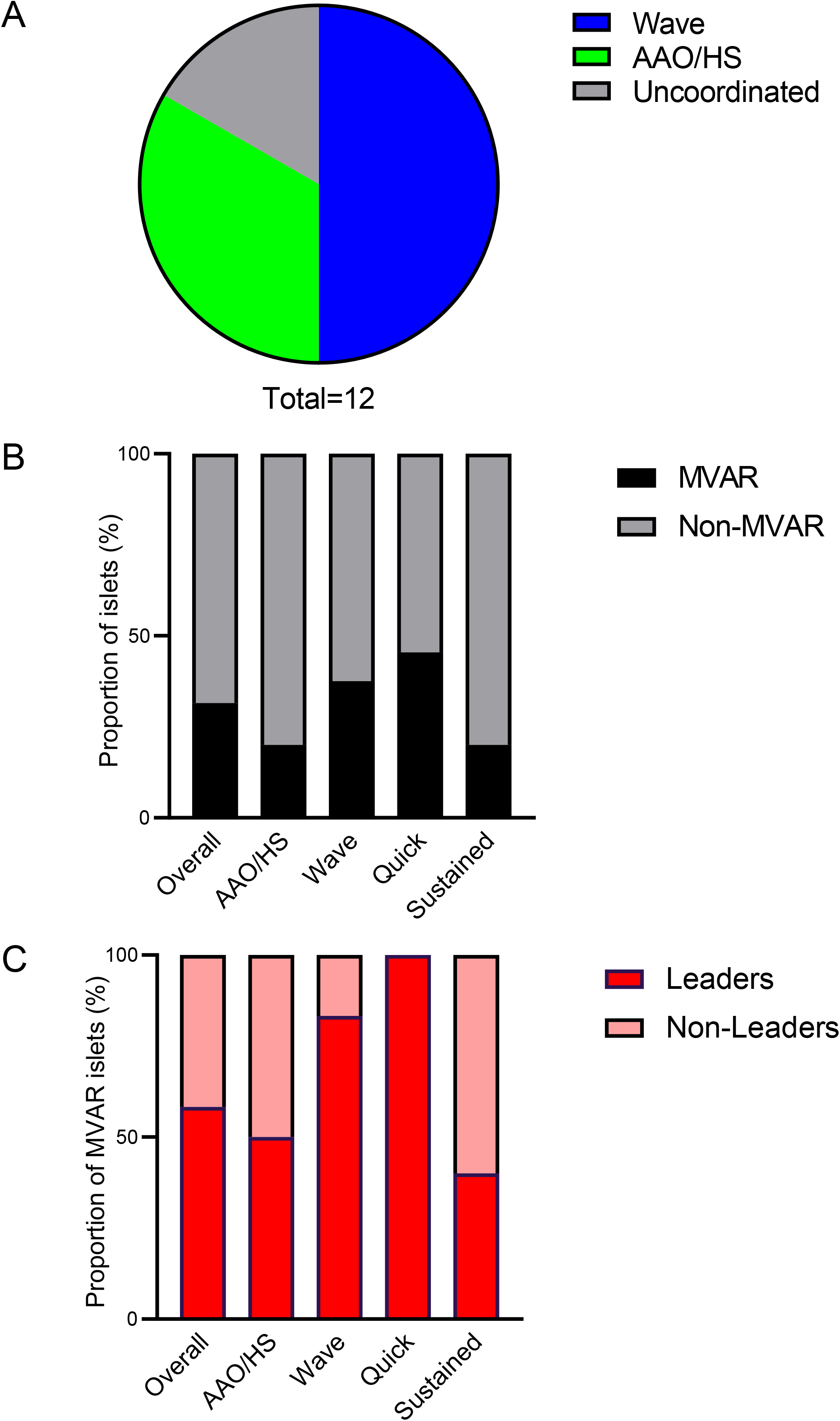
MVAR analysis. **A.** Categorisation of islets with MVAR leaders depending on signal propagation behaviour during calcium waves. 16.7% displayed uncoordinated response, 50.0% slow propagation wave >1s and 33.3% AAO high-speed waves <1s (n=12). **B.** Percentage of islets with MVAR leaders. 31.6% of all responsive islets had MVAR leaders. Each subgroup also had a proportion of islets displaying MVAR behaviour (AAO: 20.0%; Wave: 37.5%; Quick: 45.5%; Sustained: 20.0%). **C.** Percentage of MVAR islets overall and in each subgroup where leader cells were also identified to be MVAR leaders. 58.3% of islets that display MVAR behaviour had leader cells that were also identified to be MVAR leaders. High proportion of MVAR islets in each subgroup had leader cells that were also MVAR leaders (AAO: 50.0%, Wave: 83.3%, Quick: 100%, Sustained: 40.0%).

Of the islets that displayed MVAR behaviour, 58.3% had leader cells that were also identified as MVAR leaders. Subgroup analyses revealed similar proportions of islets with where leader cells were also identified to be MVAR leaders (Figure 3C; AAO: 50%; Wave: 83.3%; Quick 100%; Sustained: 40%). No highly connected or hub cells were identified as MVAR leaders.

### Leader single cell transcriptomics

We next sought to determine whether there may be stable transcriptomic differences between leader and follower cells. As described above, leaders could be identified rapidly and in real time, allowing these cells, or followers, to be labelled on the microscope stage at the end of the Ca^2+^ imaging session by photoactivation of PA-mCherry (Figure 4A). “Photopainting” was achieved by exposure to UV light at 405nm using the fluorescence recovery after photobleaching (“FRAP”) module on board the microscope (Figure 4A, B & Methods). Subsequently, islets were dissociated into individual cells which were then sorted according to fluorescence intensity (Methods). Cells which were double positive for GCaMP6 and mCherry were individually dispatched into plate wells and cDNA libraries were generated for RNA sequencing. From four fully independent experiments, we isolated a total of 14 leader cells and 9 follower (control) cells. In these experiments, 25% of islets showed quick wave behaviour vs 68% sustained, and 38% defined wave vs 56% AAO/HS; leaders or followers were not subdivided into these groups given the low numbers of cells involved.

**Figure 4.**
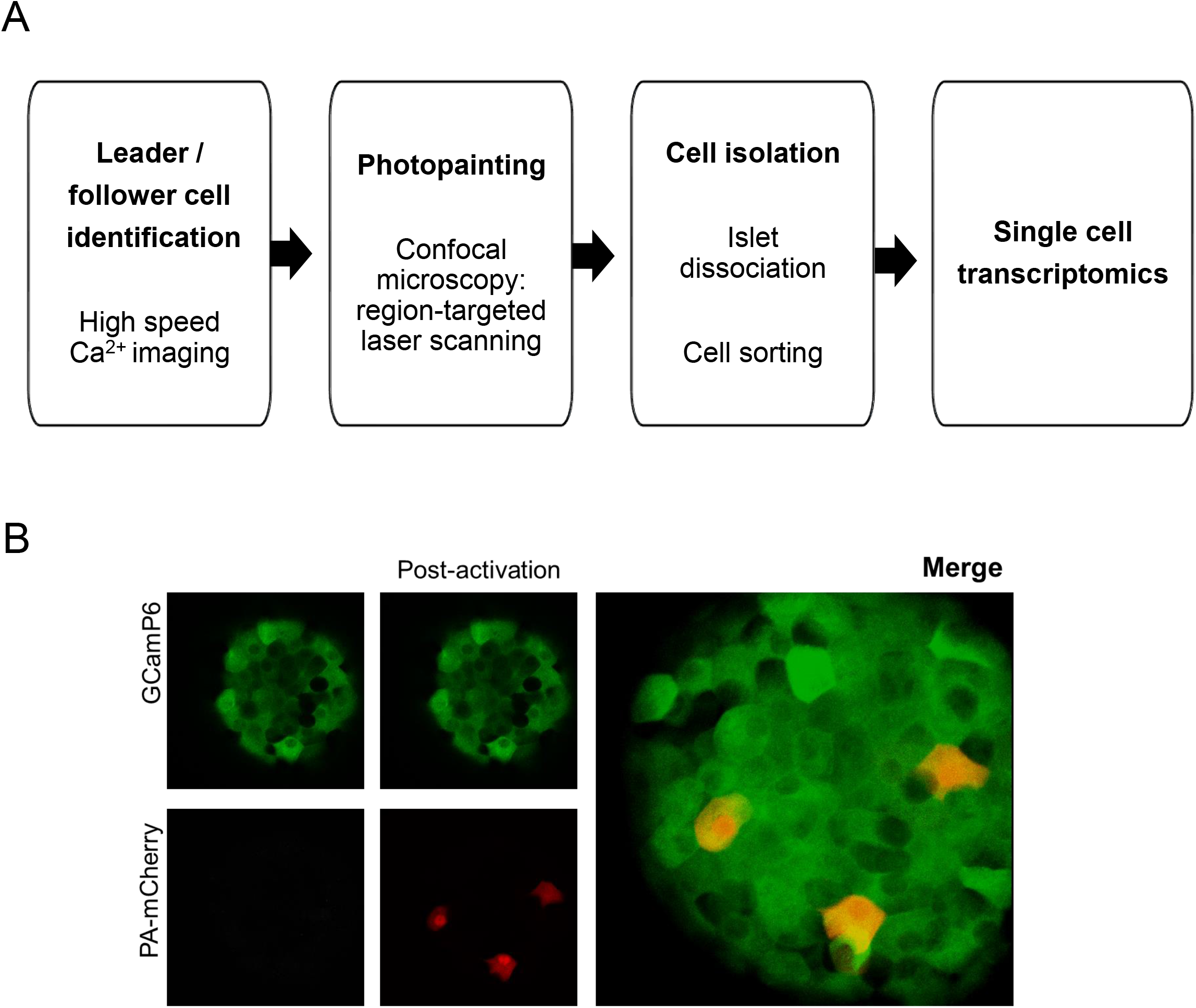
A. Experimental procedure for leader cell identification and transcriptomic analysis (“Flash-Seq”). **B.** Representative images of islet expressing GCaMP6f specifically in beta cells and infected 24 hours prior to imaging with a construct for PA-mCherry expression. Fluorescence from PA-Cherry is not detectable before exposure to UV light (left panel), while cells display high fluorescence level after targeted illumination in determined region of interest drawn around leader cells (right panel and merged image).

Sequencing data were aligned to the mouse transcriptome and differential expression (DE) analysis was performed for all transcripts (7810) that were present in at least 60% of the cells of one group (control/followers or leaders). Of these, 295 genes were differentially expressed (padj<0.05; Supplementary Table 1). Functional annotation using DAVID (Database for Annotation, Visualization and Integrated Discovery; D. W. Huang et al., 2009) revealed a substantial proportion of the differentially expressed genes to be associated with transcriptional regulation (Figure 5, Supplementary Table 2), including the most significantly upregulated gene (~180 fold, padj=0.00000002) *Bmi1*, a polycomb protein involved in beta cell proliferation [26]. Strikingly, we also identified a total of 14 genes involved in cilia function and/or assembly (Supplementary Table 2). Examples included the strongly upregulated *Adcy6, Vhl* and *Dync2i1* (~150-200 fold, padj=0.002-0.0007) and the downregulated *Dcdc2a* and *Rsph1* (~0.004 fold, padj=0.0002 and 0.01-fold, padj=0.002, respectively; Supplementary Table 1).

**Figure 5.**
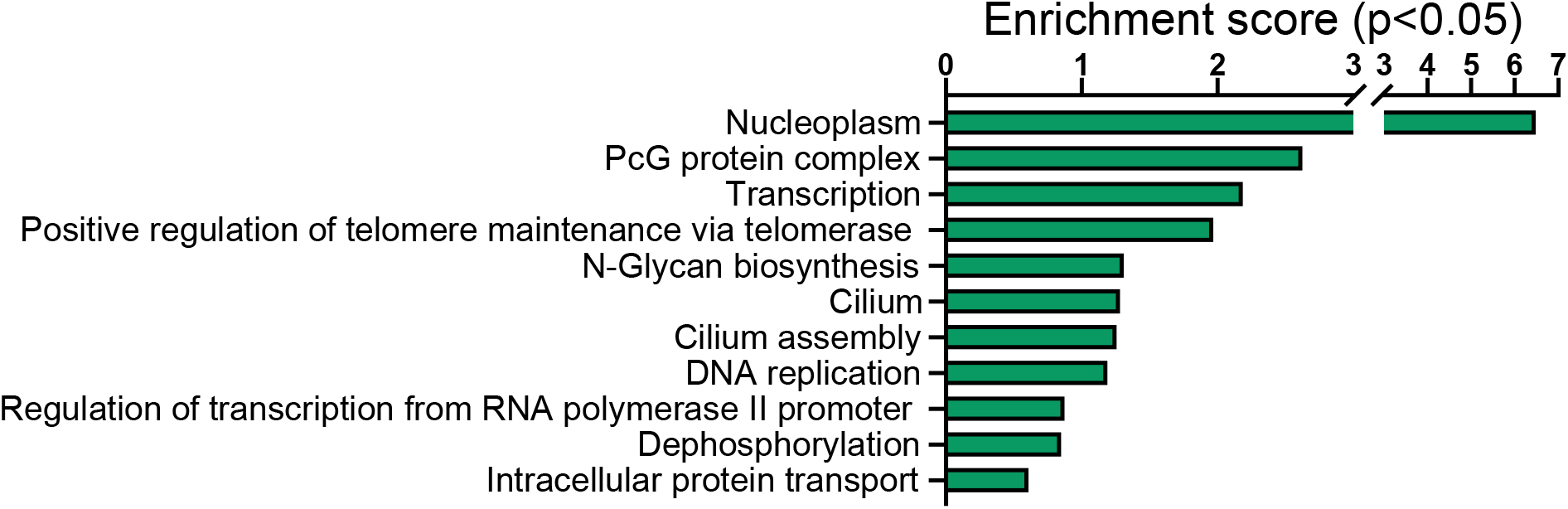
Gene ontology analysis of genes differentially expressed in leader *versus* control cells. The graph shows enrichment scores for one representative term for each cluster grouped by semantic similarities and including terms with a p value <0.05. A full list of terms is included in Supplementary Table 2.

Additionally, *Il18bp* [27] and *Cxcl16* [28] were included in the top five most significantly dysregulated genes, possibly influencing actions of cytotoxic cytokines and macrophage infiltration in the context of T1D or T2D. Also, MCUb was found among the most down-regulated gene (0.001-fold, pad=0.01). MCUb is part of the Mitochondrial Calcium Uniporter (MCU complex) and negatively regulates the activity of MCU. As such, lowered MCUb levels in leader cells may enhance calcium transfer into mitochondria, enhancing oxidative metabolism, leafing to further closure of ATP-sensitive K^+^ channels and Ca^2+^ influx [29,30].

### Gene Set Enrichment Analysis (GSEA)

We next performed GSEA as a further means of identifying additional gene ontology sets which were coordinately mis-expressed in leader versus follower cells (nominal p-values <0.05, Supplemental Table 4) In line with the functional annotation of differentially expressed genes, the top enrichment scores corresponded to gene sets associated with “Epigenetic regulation of gene expression” such as “Histone monoubiquitination”, “Euchromatin” and “Regulation of DNA template transcription elongation”, and included *BMI1* (see above). These findings are coherent with demonstrated roles for polycomb-dependent changes in the epigenome in controlling normal beta cell function [31,32].

Suggestive of a role for cell-cell interactions, we identified enrichment within the “Lateral Plasma membrane” and “Basement membrane” gene sets that included *AXIN1* and *APC*, involved in Wingless (Wnt) signalling, *LOXL2*, encoding Lysyl oxidase-like 2 involved in connective tissue remodeling and *LAMC1*, encoding laminin γ subunits.

Consistent with dysregulation of genes involved in cilium biogenesis or function (See DAVID analysis) the *GOCC_Centriolar Satellite* group was also significantly affected, and included *PARD6A* (Par6-family polarity regulator-A), also included in the *Gocc_Tight_Junction set*, and *PCNT* (pericentrin), the latter a Ca^2+^-calmodulin regulator component of the pericentrolar material, and *C2CD3* (C2-domain-containing 3 centriole elongation regulator). The *Peptide_N-Acetyl_transferase* group also included *NAA40* (N-alpha-acetyl-transferase 40) located in the centriolar satellite.

Suggesting alterations in mitochondrial metabolism, the gene set “Nucleotide Transporter” was over-represented at the bottom of the list of ranked genes and included mitochondrial solute transporters such as *SLC25A32*, involved in mitochondrial folate uptake and decreased in diabetic islets [33]. Also, in this group were *LRRC8*, Leucine-rich repeat-containing protein A, located at the plasma membrane and implicated in cell adhesion, and *ABCC5*, ABC Binding Cassette Subfamily C, member 5, involved in the export of cyclic nucleotides.

Additional affected gene sets included *Inositol Phosphate-Mediated Signalling Pathway, Smoothened Signalling Pathway* and *Response to Electrical Stimulus* (Supplementary Table 4).

### Association of mis-expressed genes in leader cells with type 2 diabetes

To determine whether any of the identified misexpressed transcripts may be associated with increased risk of Type 2 diabetes or other glycemic trait, and thus causally implicated in disease development, we accessed the T2D portal: https://t2d.hugeamp.org/. The human homologues of 215 of the identified mouse genes were located *in loci* carrying a significant (p<0.05) association with any glycemic trait (Supplementary Table 3), of which 16 displayed p-values of <10^-5^; *ADAM16, CLASRP, ERLIN1, TMEM123, CCDC186, PDE12, PARD6A, FTSJ1, FAM9C, SIDT2, CSCL16, FBX18L, TRAK2, ZNF143, BMI1, SCF8)*. One of these *(BMI1*, padj=2.2 x10^-8^) was in a locus with an association signal of <10^-7^.

We next determined whether any of the identified genes in leader cells were also misexpressed in islets in T2D. Comparison with the data sets provided by Marchetti et al. [34] revealed five common genes shared between the leader cell gene list and the organ donor set *(RSPH1, VAT1L, ZNF704, DUSP10* and *GRIA2)*, with one common gene shared with the partial pancreatectomy data set *(TMED6)*. Of the mis-expressed genes identified through single cell RNASeq by Segerstolpe et al. [11], one was common to the mouse leader gene list *(MEIS1)*.

### Leader cells tend to display lowered cilium frequency and length *versus* followers

As the transcriptome analysis pointed to a potential role of cilia biogenesis in defining leader cell characteristics, we compared the expression and morphology of primary cilia in leader *versus* follower cells (Figure 6). The cilia marker acetylated tubulin (AcTUB) was used to stain primary cilia in islets that were photopainted with mCherry for leader cells (Figure 6B, C) and beta cells were detected with green fluorescence from GCaMP6 only. Data were analysed from five independent z-projections in three islets each, selected from regions that contained clearly painted leader cells. A total of 118 leader cells and 980 follower cells were identified in these stacks and were examined for ciliation status and ciliary length. A range of 41.2% to 91.5% (average 67.6%) of leader cells and a range of 29.8% to 94.6% (average 74.0%) of follower cells contained cilia (Figure 6D). Cilia length averaged 3.76 ±0.18μm and 4.08±0.07μm in leader and follower cells respectively (Figure 6E). Non–significant tendencies were observed for lower cilium frequency (p=0.098; Figure 6D) and length (p=0.142; Figure 6E) in leader *versus* follower cells.

**Figure 6.**
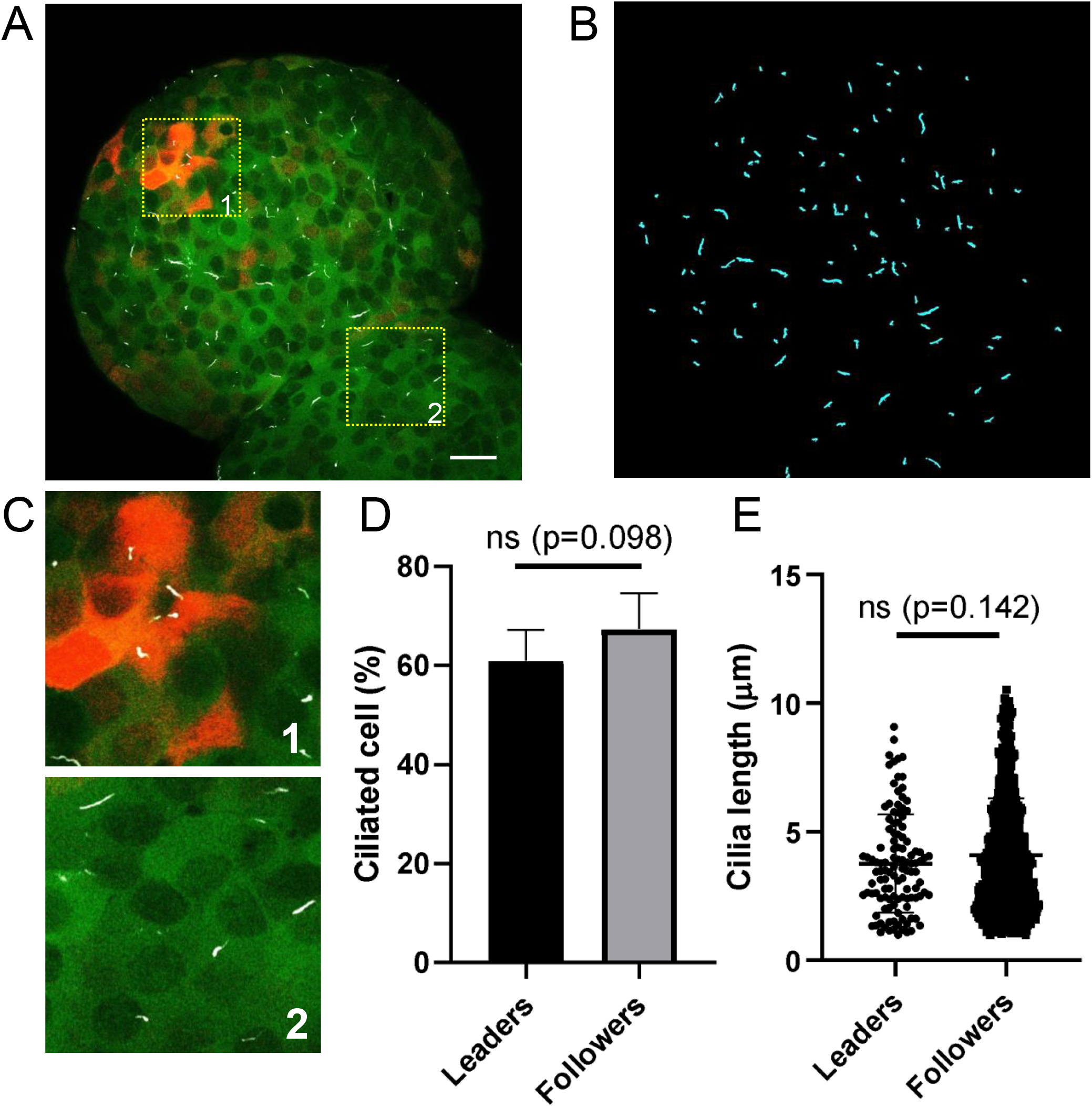
Quantification of cilium frequency and length. **A.** Representative islet crosssection labelled with primary cilia (white), PA-mCherry leader cells (red), and beta cell GCaMP6f (green). Scale 10 μm. **B.** Segmented cilia after maximum intensity projection of islet slice and manual correction in CiliaQ Preparator and Editor. **C.** Insets showing representative regions containing (1) leader cells labeled in both green and red, (2) follower cells which are green only. Both leader and follower cells are ciliated (white). **D.** Cilia frequency quantitation showing comparable levels of ciliation among leader and follower cells, p = 0.098. **E.** Cilia length quantitation showing a trend toward shorter cilia in leader cells that did not reach statistically significance, p = 0.142.

### Leader cells are located closer to delta cells than are followers

It has been suggested that paracrine regulation by delta cells may contribute to, or restrict, the transmission of Ca^2+^ waves between beta cells within the islet [35,36]. To explore this possibility, we measured the 3D Euclidian distances between labelled leaders (PA-mCherry) and follower beta cells (GCaMP6^+^) to delta cells (somatostatin+) *post hoc* (Methods, Figure 7). The average distance from the photo-labelled leader cells to the nearest delta cell was 18.2 ± 9.1 μm *(n*=15 leader cells, 443 somatostatin+ cells from 6 islets). In contrast, the average distance of follower cells to delta cells was 61.8±15.4 μm *(n*=5127 follower cells, 443 somatostatin+ cells from 6 islets; paired two-tailed Student’s t-test, p<0.002).

**Figure 7.**
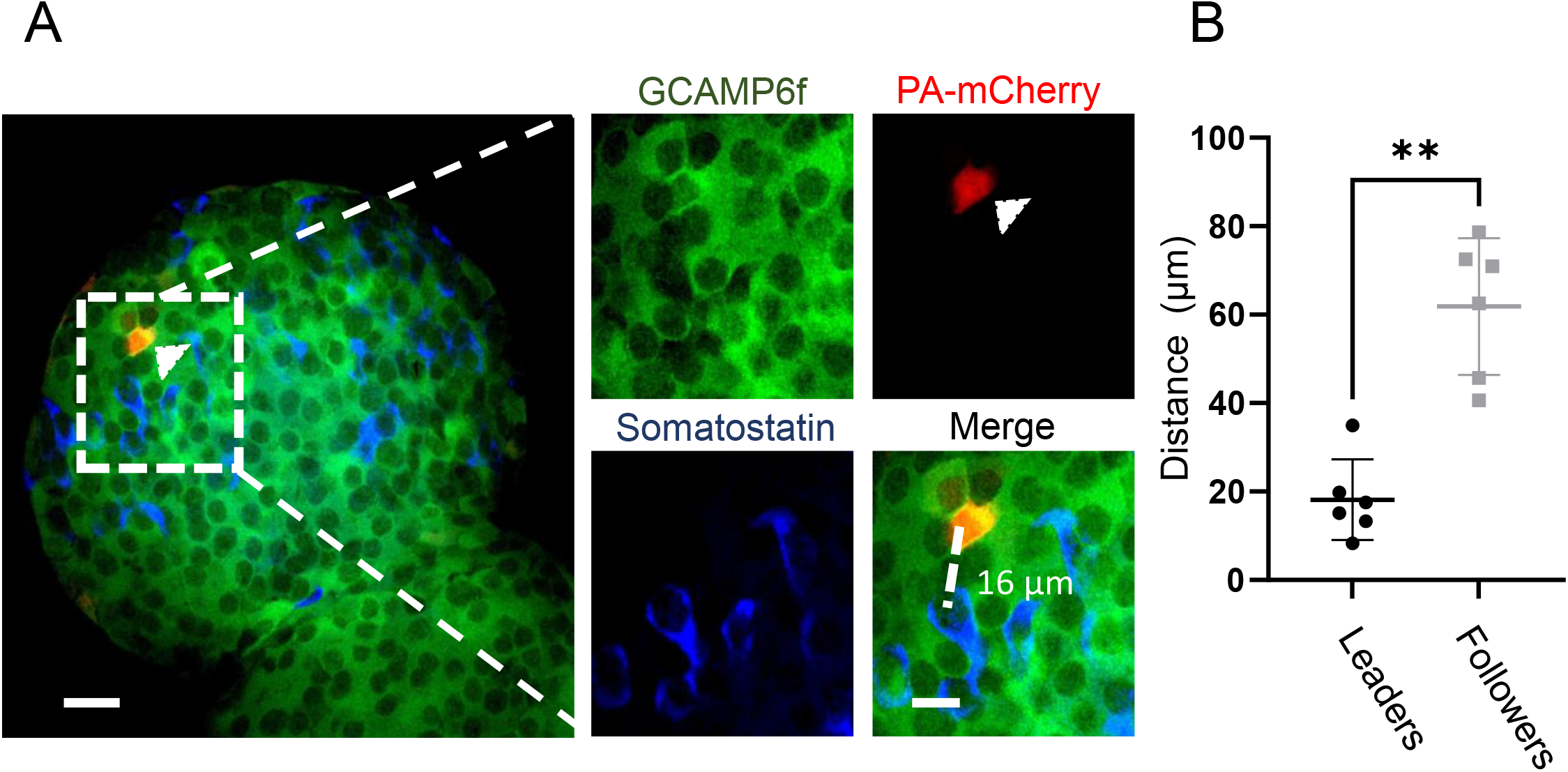
3D Euclidian distance of leader and follower beta cells towards SST+ cells. **A.** Confocal image of the photo-painted islet. GCaMP6f (green), somatostatin (blue), and photo-labelled PA-mCherry (red) mark the beta cells, the delta cells and a leader beta cell, respectively. **B**. Quantification of the 3D Euclidian distance from leader beta cells and follower beta cells to delta cells islets from C57BL/6J Ins1Cre:GCaMP6f^fl/fl^ transgenic mouse. Scale bars: 20 μm. See the text for further details.

## DISCUSSION

The overall aims of the current study were, firstly, to identify and characterize beta cell subpopulations with discrete roles in Ca^2+^ wave initiation and propagation across the islet. Secondly, we sought to determine whether differences may exist at the transcriptomic level between leader and follower cells, and between the localisation of leaders *vs* followers vis a vis their proximity to delta cells. Whereas the latter are stimulated by elevated glucose concentrations and may conceivably play a paracrine role in controlling beta cell behaviour, alpha cells are usually inactivated at high glucose (at least in rodent islets).

An important finding was that only ~1/3 of islets display small worlds topography, and the existence of highly connected hubs, as described by ourselves and others in recent years [16,17,37]. This may reflect differences between the methodologies used in the present and earlier studies. For example, we used here the fluorescent protein based Ca^2+^ sensor GCaMP6 rather than a low molecular weight chemical probe such as Fluo2. Intracellular buffering is likely to affect calcium dynamics, and the greater effective concentration of calcium binding sites when using low molecular weight probes may thus dampen fluctuations in Ca^2+^. Indeed, we observed more wave-like behaviour emanating from well-defined leader or groups of leader cells in recent studies in which GCaMP6-expressing islets were engrafted into the anterior eye chamber of the eye [17,38], versus studies using low molecular weight probes in isolated islets [16,39], though hub-leader dynamics were nevertheless observed in the eye chamber setting. However, we cannot exclude the possibility that there are actions of the calmodulin moiety present in GCaMP6 on calcium-regulated proteins and processes in GCaMP6-expressing cells. Consistent with the concentration of calcium-binding sites being a key determinant in differing calcium dynamics, our preliminary data (not shown), as well as those of others [40], using the bright Cal520 probe demonstrate a greater propensity towards wave-like, and less small-worlds behaviour. Importantly, we observed almost no overlap between leader cells and highly connected hub cells. Similar conclusions were recently reached by Kravets and colleagues [24].

A further unexpected finding was the relatively small proportion of islets that had MVAR leaders. MVAR includes Granger causality analysis, as used in our previous study [17], and both seek to identify cells whose behaviour predicts that of other cells in the islet. Hence, MVAR leaders are defined as a subset of beta cells that, mathematically, cause other beta cells to respond. It should be emphasised that even though MVAR leaders were only found in a third of the responsive islets, causal relationships were still found between beta cells in the other 68.4% of islets. However, these islets were not considered to house MVAR leaders because no observable hierarchy of beta cell response was present. Of note, MVAR leaders were found across all four subgroups, indicating that small-worlds behaviour was not necessary for their presence.

We also demonstrate in the present study the existence of a “main” leader cell, alongside other leaders associated with a smaller number of Ca^2+^ waves. Thus, and although a few cells (on average 2-3/islet optical section) could be observed during the time of acquisition, a single cell was at the origin of most of the oscillations (~60%) while a secondary leader controlled ~30%, and 1 – 2 leader cells the minority of the remaining oscillations. This indicates that leaders themselves represent a heterogeneous subpopulation. Interestingly, in mouse islets, the presence of more than one leader cells leads us to hypothesize that there may be partial redundancy in leader cell function, wherein inactivation of the “main” leader cell could result in a secondary or tertiary leader taking over. Whether follower cells also play an active role as suppressors of Ca^2+^ oscillation initiation remains to be explored, though our previous studies [16] did not reveal any increase in Ca^2+^ oscillations when these cells (rather than hubs) were temporally inactivated by optogene stimulation.

We have developed and used here a novel strategy, which we refer to as “Flash-Seq”, to interrogate the transcriptomes of leader or follower cells selected according to our analysis of calcium dynamics, and then selectively labelled by photo-activation of the fluorescent probe, PA-mCherry. Inherently with the experimental methods, the number cells collected was very low, as the identification of leader cells only depends on individual cells identification after high-speed calcium imaging. Whilst this approach highlighted the existence of a substantial number (~300) of significantly miss-expressed genes in leader versus the follower population, we would emphasise several challenges in the analysis of these data obtained from a relatively small number (23) of individual cells. Firstly, we noted that PCA analysis did not provide a clear separation between the groups (leader vs follower). Secondly, we noticed that transcript levels showed frequent drop-outs, which resulted in undetectable expression of a specific transcript in a minority of cells, even when other cells in the same experiment and group (leader versus follower) had extremely high levels of expression. The drop-out rate was not influenced by the total number of transcripts expressed in the cell or other characteristics of the RNA-seq datasets. This phenomenon was consistent with other single cell transcriptome studies [41], where it has usually been dealt with by comparing gene expression between groups with over several hundred or thousand clustered single cells or by imputation methods [42,43]. Since neither approach was compatible with our data, we chose instead a “simple” filtering approach to account for sparsity in the data, which consisted in excluding those genes that were not detected in at least 60% of the samples of one of the groups from the differential expression analysis.

Perhaps not surprisingly, a substantial proportion of the miss-regulated genes are involved in transcriptional control, and thus potentially in the control of cellular identity. These included several members of the polycomb repressor complex, such as *Bmi1*. Polycomb proteins are well-established chromatin modifiers and thus our data supports a role for chromatin remodelling in the establishment/maintenance of leader and follower cells within the islets. Interestingly, intra-islet beta cell heterogeneity in polycomb states has already been reported, and beta-cell polycomb loss of function triggers diabetes-like transcriptional signatures and de-differentiation [31]. It is tempting to hypothesise that an alteration of the leader-follower roles within these islets may contribute to these defects. *Bmi1* plays a positive role in proliferation during beta cell development by restricting p16^ink4^ expression [26]. Nevertheless, *Cdkn2a* (encoding p16^ink4^) expression is similar in leader and follower cells, indicating a p16^ink4^-independent effects of *Bmi1* in defining these populations. Of note, the human *BMI1* gene lies in a locus quite strongly (padj <10^-7^) associated with glycemic risk, suggesting that inter-individual differences in *BMI1* expression in beta cells may influence leader-follower dynamics to impair normal insulin secretion.

Of note, expression of neither insulin nor glucokinase, previously shown to be differentially expressed at the protein level between hub and follower cells [16,17], were impacted in leader cells, in accordance with the view that these cells also do not overlap substantially with hub cells.

Remarkably, a significant number of genes associated with cilium biogenesis and assembly cilia were represented in the filtered list of differentially-expressed genes. Primary cilia are sensory organelles on the surface of islet alpha, beta, and delta cells and play key roles in regulating hormone secretion and cell-cell communication [44–46]. Recently, beta cell cilia have been reported to be motile as a dynamic regulator of beta cell calcium and insulin response to glucose [47], and genes modulating cilia biogenesis and remodelling have been found enriched in human T2D beta cells [48]. The primary cilium has previously been shown to maintain a functional beta cell population, and ciliary dysfunction impairs beta-cell insulin secretion and promotes development of type 2 diabetes in rodents [49,50]. We now find using FLASH-Seq that leader cells are defined by differential expression of cilia and centrosome structural components, signalling factors, and ciliogenesis regulators (Supplemental Table 2). These include centriolar assembly factors CCDC, pericentrin, signalling factors adenylyl cyclase and Rab, motile cilia genes dynein and radial spoke head component 1, and known human ciliopathy genes such as *Tmem138* and *Bbs1* which are linked to Joubert and Bardet-Biedl syndromes. Our first-pass analysis revealed clear tendencies towards lowered cilia frequency and length in leader cells. It is equally possible that cilia differences might go beyond static morphological measures. In particular, our discovery of motile cilia genes in leader beta cells supports the recently reported phenomenon of beta cell cilia motility, which modulates calcium and insulin secretory responses to glucose [47]. We speculate that, beyond static structure, primary cilia on leader cells may exhibit differential capacity for movement, nutrient sensing, or signalling, giving these cells the ability to initiate Ca^2+^ waves early. Examination of live-cell cilia dynamics in leader beta cells therefore would be an important future experiment. In conclusion, despite the limitations of our approach as discussed above, the present results provide compelling evidence for the existence of leader cells with a discrete molecular phenotype from follower cells. Future experiments, involving larger numbers of cells, should allow these transcriptomic differences to be refined and extended, and ultimately to be supported by analyses at the level of the proteome. The importance of the mis-expressed genes may in the future be revealed by studies in which the expression of these is modified across the beta cell population or, ideally, at the level of individual cells. Nevertheless, an important conclusion from the present studies is that the behaviours described for discrete subpopulations are unlikely to result purely from of the localisation of these cells within the islet, including proximity to other islet cell types, nerve endings, blood vessels and so on. Nevertheless, we do not exclude a role for these parameters. Indeed, and strikingly, we show that leaders are located substantially closer to delta cells than are followers. Although this result may appear paradoxical insofar as somatostatin secretion from delta cells is expected to exert an inhibitory effect on neighbouring beta cells, it is conceivable that the lower number and length of cilia – where somatostatin receptors including SSTR3 [51–54] are concentrated – may lead to a complex interplay between the leader cell and nearest beta cell neighbours which influence the initiation of Ca^2+^ increases. This may provide an example of how a discrete molecular phenotype may interact with a defined localisation to purpose leader cells for their role.

## METHODS

### PA-mCherry adenovirus construct generation

PA-mCherry1-C1 plasmid was from Addgene (gift from Dr. Michael Davidson Addgene plasmid # 54495; http://n2t.net/addgene:54495; RRID:Addgene_54495). The PA-mCherry sequence was cloned into pShuttle-CMV plasmid (gift from Dr. Bert Vogelstein – Addgene plasmid # 16403; http://n2t.net/addgene:16403; RRID:Addgene_16403). An adenovirus construct was generated using the pAdeasy system[55].

### Ca^2+^ imaging in mouse islets

Colonies of Ins1Cre.GCCaMP6^f/f^ mice were maintained on a C57/BL6 background on regular chow and under controlled temperature (21 – 23 °C), humidity (45 – 50%) and light (12:12 h light-dark schedule, lights on at 0700 hours; Salem et al., 2019). Local ethical committee approval was obtained (Imperial College AWERB; CRCHUM, Montreal CIPA 2022-10040 CM21022GRs). Experiments in the U.K were performed under Home Office License PA03F7F0F (IL).

Islets were isolated from male mice aged 8-12 weeks, as previously described [56]. In brief, pancreata were inflated by injecting collagenase-containing media (Sigma-Aldrich; 1mg.mL^-1^) into the pancreatic duct, followed by pancreas isolation, exocrine tissue digestion for 10 minutes at 37 °C and islets separation by gradient density [56]. GCamP6f-expressing islets were then cultured in RPMI 1640 medium (GIBCO) containing 11 mM glucose supplemented with 2 mM L-glutamine, 100 IU/mL penicillin, and 100 μg/mL streptomycin (GIBCO) and foetal bovine serum (FBS – 10% v/v) at 37 °C in 5% CO_2_ humidified atmosphere. Islets were infected with PA-mCherry adenovirus construct (100 Multiplicity of Infection) 24h post isolation and Ca^2+^ imaging was performed 24 h post infection. Islets were transferred from culture media into an imaging chamber containing Krebs-HEPES-bicarbonate (KHB) buffer (130mM NaCl, 3.6mM KCl, 1.5mM CaCl_2_, 0.5mM MgSO_4_, 0.5mM NaH_2_PO_4_, 24mM NaHCO_3_, 10mM HEPES; pH 7.4) complemented with 11 mM glucose and imaged on a Zeiss LSM-780 inverted confocal microscope equipped notably with an incubation system for temperature control (37°C), laser lines including 405, 488 and 561 nm and a 20x/0.8 Plan achromat objective. Acquisitions were performed at a ratio of 5 images.s^-1^ on single plane (256×256 pixels – 415×415 μm) without averaging, using the 488nm laser line for GCamP6f excitation, for a total time of 10 min. High-speed calcium imaging acquisitions were later analysed for connectivity analysis as described below. Raster plots were generated off-line based on the approach of Rupnik and colleagues [57] using Python scripts provided at https://github.com/szarma/Physio_Ca.

### Photopainting and isolation of leader or follower cells

At the end of each high-speed calcium acquisition, movies were screened to identify leader cells, defined as the first cell to display a Ca^2+^ signal increase (>20% of baseline) during an oscillation in response to 11mM glucose – i.e. a global activation of the islets, with >75% of cells showing a Ca^2+^ signal increase of >20% of baseline (Figure 1A-D). When possible, a leader cell was identified for each oscillation occurring during the 10 min acquisition period. We used the FRAP module in the acquisition software ZEN controlling the LSM780 microscope, which allows to illuminate with the scanning laser a specific region of interest (ROI) drawn on a previously acquired image. Thus, after calcium imaging, a ROI was drawn around each leader cells, that were then specifically scanned using the 405nm laser line at 50% for 3 occurrences, to elicit PA-mCherry photoactivation. Correct photoactivation in individual leader cells was assessed by imaging using 561 nm laser before and after UV illumination (Figure 3B).

Post photoactivation, islets were put back to culture and on average 10 to 15 islets were individually processed per experiment (Figure 3A). Dissociation was performed by incubating islets in accutase (GIBCO, 50 islets in 500μL) for 5 min at 37°C and mechanically disaggregated by trituration. After adding 1mL of culture media to accutase solution, dissociated cells were pelleted, washed in PBS and resuspended in cell sorting buffer at 4°C (PBS pH7.4, FBS 2% v/v, 1mM EDTA). Cells were then sorted at 4°C on a fluorescence-activated cell sorters BD Aria Fusion with settings selecting cells with both high fluorescence signals for GCamP6f (excitation 488nm) and for mCherry (excitation 561nm). Single cells were dispatched into a 96-plate well containing 5μL 1x lysis buffer containing Murine RNase inhibitor (NEBNext Single Cell/Low Input RNA Library Prep Kit for Illumina).

Control cells were selected using the same protocol where follower cells were photoactivated.

### Transcriptomics analysis

cDNA libraries were generated from 14 leader and 9 follower single cells using the NEBNext Single Cell/Low Input RNA Library Prep Kit for Illumina according to the furnisher protocol. Briefly, after single cells were lysed in 5μL 1x lysis buffer (see above), reverse transcription and template switching steps were performed, and obtained cDNA was amplified by PCR (20 cycles). After clean-up, cDNA quality and quantity were tested with a Bioanalyzer (Agilent) using DNA High Sensitivity chips. Then, a cDNA fragmentation step was performed followed by ends preparation for adaptor ligation (NEBNext Adaptor for Illumina; 0.3 μM). After cleanup of adaptor-ligated DNA, a PCR enrichment step was performed (8 cycles) using unique primer pairs for each library (NEBNext Multiplex Oligos for Illumina-Dual Index Primers Set 1).

Library quality was assessed with a Bioanalyzer High Sensitivity chip, and an average fragment size of 300 pb was confirmed before sequencing in an Illumina HiSeq 4000. 3.8-5.4 million reads per cell were successfully aligned to the mouse transcriptome (GenCode M23) with Salmon v1.3 [58]. Differential expression analysis performed with DESeq2 v1.28.0 in R v4.0.2 [59]. Only transcripts that were present in at least 60% of the samples of one of the two groups (leaders or followers) where included in the analysis. Differentially expressed genes (padj<0.05) were subjected to Gene Ontology analysis in the Database for Annotation, Visualization and Integrated Discovery, DAVID [25].

### Connectivity analysis

#### Signal binarisation

Ca^2+^ signals were denoised by subjecting the signal to the Huang-Hilbert type (HHT) empirical mode decomposition (EMD), as used in previous studies [16,17]. The signals were decomposed into their intrinsic mode functions (IMFs) in MATLAB [60]. The residual and the first IMF with the high-frequency components were then rejected to remove random noise.

The Hilbert-Huang Transform was then performed to retrieve the instantaneous frequencies [61–63] of the other IMFs to reconstruct the new signal using

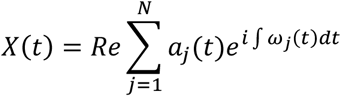

where *a_j_*(*t*) = amplitude, *ω_j_*(*t*) = frequency of the *i*th IMF component [64] to retrieve a baseline trend and to account for any photobleaching or movement artefacts. A 20% threshold was imposed to minimise false positives from any residual fluctuations in baseline fluorescence.

Cell signals with deflection above the de-trended baseline were represented as ‘1’ and inactivity represented as ‘0’, thus binarising the signal at each time point. The coactivity of every cell pair was then measured as:

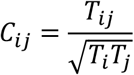

where *T_ij_* = total coactivity time, *T_i_* and *T_j_* = total activity time for two cells

The significance at p <.001 of each coactivity measured against chance was assessed by subjecting the activity events of each cell to a Monte Carlo simulation [65,66] with 10,000 iterations.

Synchronised Ca^2+^-spiking behaviour was assessed by calculating the percentage of coactivity using the binarised cell activity dataset. A topographic representation of the connectivity was plotted in MATLAB [67] with the edge colours representing the strength of the coactivity between any two cells.

A 80% threshold was imposed to determine the probability of the data, which was then plotted as a function of the number of connections for each cell to determine if the dataset obeyed a power-law relationship [68,69].

Finally, the data and figures were written from MATLAB to Microsoft Excel files [70] for easy visualisation and dissemination.

#### Multivariate vector autoregression

Multivariate vector autoregression (MVAR) is a stochastic process model that captures any linear interdependencies in a time series while allowing for dynamic multivariate time series modelling [71,72]. Its theory is based on the Granger causality test [73,74], a statistical hypothesis used in previous studies [17] to examine causality between beta cells in the islets.

A vector autoregressive model was built in Jupyter Notebook (Python 3) based on:

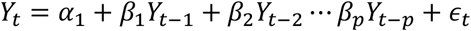

where α = intercept, β1 and β2 till βp = coefficients of the lags of Y till order p

First, the time series data of all the signals were visually analysed for any outstanding characteristics. Next, causality amongst all time series was tested using first the Granger’s Causality Test to prove that the past values of one time series have a causal effect on another series [71,73,75–77]. Following that, cointegration of the data was tested to establish the presence of any statistically significant connection between two or more time series [76–78].

Once it was established that there was a statistically significant connection and causation amongst the time series data, stationarity was checked using the Augmented Dickey Fuller Test [79] to ensure that the mean and variance of every time series did not change over time. Any non-stationary series was made stationary by differentiating all series until all the time series data reach stationarity.

Finally, the data and figures were written from Jupyter Notebook to Microsoft Excel files for easy visualisation and dissemination.

#### Gene list overlapping

Of a total of 295 mouse genes that showed differentially expressed (adjusted p-value<0.5) between the leader and the follower cells, 268 genes were identified and converted to their human equivalent. The r package babelgene was used for converting between human and non-human gene orthologs/homologs, which is sourced from multiple datasets and compiled by the HGNC Comparison of Orthology Predictions (HCOP) [80–82]. In the type 2 diabetes knowledge portal (https://t2d.hugeamp.org/), our dysregulated gene list was then compared with a list of genes that are either the closest genes to a lead SNP which shows associations (p-value ≤ 5e-8) for glycemic phenotype, or the genes with a significant (p-value ≤ 2.5e-6) gene-level association for the glycemic phenotype (Multi-marker Analysis of GenoMic Annotation (MAGMA) method.

#### Euclidian distance between cells

After images were acquired, the nuclei of each cell labelled by GCaMP6f(beta-cells), SST (delta-cells) and photo-labelled PA-mCherry (leader beta-cell), the 3D Euclidian distance was calculated [83]. Briefly images were loaded in ImageJ, the center of the nuclei was selected manually, and the X, Y and Z coordinates for each nucleus was extracted. To calculate the distance, we used the following formula:

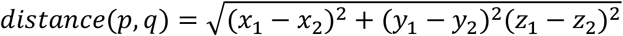

where *p* and *q* being the center for each cell while *x, y, z* are the X, Y and Z coordinates for each nucleus.

#### Cilia imaging and quantitation

Photopainted mouse islets were fixed in 4% paraformaldehyde (PFA) and permeabilized with 0.3% Triton X-100 in PBS (PBST) prior to blocking and staining with rabbit anti-acetylated alpha tubulin antibody (Cell Signaling #5335). Confocal 3D images of whole islets were used for cilia frequency and length quantitation. Individual 3 to 5μm-thick optical slices with 1μm step size were selected from lower, mid, and upper regions of the islets, each slice spanning a single layer of cell body and cilia. 14 individual slices were evaluated from 3 islets. In total, analysis was performed on 118 leader cells and 980 follower cells. Cilia frequency and length were measured using ImageJ with CiliaQ v0.1.4 plugins [84]. Images were first processed and segmented with CiliaQ Preparator using Canny 3D edge detection in maximum intensity projections with low threshold delivered by algorithm Triangle and high threshold by algorithm Otsu, Gaussian Blur with sigma 1.0. After segmenting the cilia from background in the acetylated tubulin channel, pre-processed images were annotated and edited with CiliaQ Editor. Extensive review and editing were required for certain islet regions such as the periphery which contained nerves and cell bodies that also stained strongly for acetylated tubulin. In these cases, manual correction was required to remove non-ciliary structures and to trace or exclude any incompletely segmented cilia. Final processed images were analysed using CiliaQ. Cilia frequency was calculated by dividing the number of ciliated cells by total number of cells from each segment. Statistical significance was determined using two-tailed t-test assuming equal variances. Cilia <1μm in length or which were abnormally long due to image overlap or possible actual fusion were excluded from the final analysis.

#### Statistics

Data are expressed as mean±SEM unless otherwise stated. Significance was tested by Student’s two-tailed t-test in MATLAB R2020a, with p<0.001 being considered significant. Significance was also tested by one-way ANOVA with Tukey’s Honestly Significant Difference (HSD) multiple comparison test for comparison of more than two groups, using MATLAB R2020a. p<0.05 was considered significant.

#### Data availability

RNA Seq data will be deposited in a publicly available repository (Gene Expression Omnibus) upon acceptance. MATLAB and Python scripts will be available on Github upon acceptance.

## Acknowledgements

We thank Samuel Reyd and Virgile Rettault for help in the generation of Raster plots. We thank the LMS/NIHR Imperial Biomedical Research Centre Flow Cytometry Facility for support.

## REFERENCES

[1] N. Esser, K.M. Utzschneider, S.E. Kahn, Early beta cell dysfunction vs insulin hypersecretion as the primary event in the pathogenesis of dysglycaemia, Diabetologia. 63 (2020) 2007–2021. https://doi.org/10.1007/s00125-020-05245-x.

[2] D.J. Steiner, A. Kim, K. Miller, M. Hara, Pancreatic islet plasticity: interspecies comparison of islet architecture and composition, Islets. 2 (2010) 135–145. https://doi.org/10.4161/isl.2.3.11815.

[3] D. Salomon, P. Meda, Heterogeneity and contact-dependent regulation of hormone secretion by individual B cells, Exp. Cell Res. 162 (1986) 507–520. https://doi.org/10.1016/0014-4827(86)90354-x.

[4] D. Bosco, P. Meda, Actively synthesizing beta-cells secrete preferentially after glucose stimulation, Endocrinology. 129 (1991) 3157–3166. https://doi.org/10.1210/endo-129-6-3157.

[5] R. Kiekens, P. In ‘t Veld, T. Mahler, F. Schuit, M. Van De Winkel, D. Pipeleers, Differences in glucose recognition by individual rat pancreatic B cells are associated with intercellular differences in glucose-induced biosynthetic activity, J. Clin. Invest. 89 (1992) 117–125. https://doi.org/10.1172/JCI115551.

[6] C.F. Van Schravendijk, R. Kiekens, D.G. Pipeleers, Pancreatic beta cell heterogeneity in glucose-induced insulin secretion, J. Biol. Chem. 267 (1992) 21344–21348.

[7] A. Wojtusciszyn, M. Armanet, P. Morel, T. Berney, D. Bosco, Insulin secretion from human beta cells is heterogeneous and dependent on cell-to-cell contacts, Diabetologia. 51 (2008) 1843–1852. https://doi.org/10.1007/s00125-008-1103-z.

[8] A.C. Carrano, F. Mulas, C. Zeng, M. Sander, Interrogating islets in health and disease with singlecell technologies, Mol. Metab. 6 (2017) 991–1001. https://doi.org/10.1016/j.molmet.2017.04.012.

[9] J. Li, J. Klughammer, M. Farlik, T. Penz, A. Spittler, C. Barbieux, E. Berishvili, C. Bock, S. Kubicek, Single-cell transcriptomes reveal characteristic features of human pancreatic islet cell types, EMBO Rep. 17 (2016) 178–187. https://doi.org/10.15252/embr.201540946.

[10] R.B. Prasad, L. Groop, Single-Cell Sequencing of Human Pancreatic Islets-New Kids on the Block, Cell Metab. 24 (2016) 523–524. https://doi.org/10.1016/j.cmet.2016.09.012.

[11] Å. Segerstolpe, A. Palasantza, P. Eliasson, E.-M. Andersson, A.-C. Andréasson, X. Sun, S. Picelli, A. Sabirsh, M. Clausen, M.K. Bjursell, D.M. Smith, M. Kasper, C. Ämmälä, R. Sandberg, Single-Cell Transcriptome Profiling of Human Pancreatic Islets in Health and Type 2 Diabetes, Cell Metab. 24 (2016) 593–607. https://doi.org/10.1016/j.cmet.2016.08.020.

[12] Y.J. Wang, K.H. Kaestner, Single-Cell RNA-Seq of the Pancreatic Islets--a Promise Not yet Fulfilled?, Cell Metab. 29 (2019) 539–544. https://doi.org/10.1016/j.cmet.2018.11.016.

[13] Y.J. Wang, J. Schug, K.-J. Won, C. Liu, A. Naji, D. Avrahami, M.L. Golson, K.H. Kaestner, Single-Cell Transcriptomics of the Human Endocrine Pancreas, Diabetes. 65 (2016) 3028–3038. https://doi.org/10.2337/db16-0405.

[14] Y. Xin, J. Kim, H. Okamoto, M. Ni, Y. Wei, C. Adler, A.J. Murphy, G.D. Yancopoulos, C. Lin, J. Gromada, RNA Sequencing of Single Human Islet Cells Reveals Type 2 Diabetes Genes, Cell Metab. 24 (2016) 608–615. https://doi.org/10.1016/j.cmet.2016.08.018.

[15] E. Bader, A. Migliorini, M. Gegg, N. Moruzzi, J. Gerdes, S.S. Roscioni, M. Bakhti, E. Brandl, M. Irmler, J. Beckers, M. Aichler, A. Feuchtinger, C. Leitzinger, H. Zischka, R. Wang-Sattler, M. Jastroch, M. Tschöp, F. Machicao, H. Staiger, H.-U. Häring, H. Chmelova, J.A. Chouinard, N. Oskolkov, O. Korsgren, S. Speier, H. Lickert, Identification of proliferative and mature β-cells in the islets of Langerhans, Nature. 535 (2016) 430–434. https://doi.org/10.1038/nature18624.

[16] N.R. Johnston, R.K. Mitchell, E. Haythorne, M.P. Pessoa, F. Semplici, J. Ferrer, L. Piemonti, P. Marchetti, M. Bugliani, D. Bosco, E. Berishvili, P. Duncanson, M. Watkinson, J. Broichhagen, D. Trauner, G.A. Rutter, D.J. Hodson, Beta Cell Hubs Dictate Pancreatic Islet Responses to Glucose, Cell Metab. 24 (2016) 389–401. https://doi.org/10.1016/j.cmet.2016.06.020.

[17] V. Salem, L.D. Silva, K. Suba, E. Georgiadou, S. Neda Mousavy Gharavy, N. Akhtar, A. Martin-Alonso, D.C.A. Gaboriau, S.M. Rothery, T. Stylianides, G. Carrat, T.J. Pullen, S.P. Singh, D.J. Hodson, I. Leclerc, A.M.J. Shapiro, P. Marchetti, L.J.B. Briant, W. Distaso, N. Ninov, G.A. Rutter, Leader β-cells coordinate Ca2+ dynamics across pancreatic islets in vivo, Nat. Metab. 1 (2019) 615–629. https://doi.org/10.1038/s42255-019-0075-2.

[18] M. Gosak, A. Stožer, R. Markovič, J. Dolenšek, M. Perc, M.S. Rupnik, M. Marhl, Critical and Supercritical Spatiotemporal Calcium Dynamics in Beta Cells, Front. Physiol. 8 (2017) 1106. https://doi.org/10.3389/fphys.2017.01106.

[19] M.J. Westacott, N.W.F. Ludin, R.K.P. Benninger, Spatially Organized β-Cell Subpopulations Control Electrical Dynamics across Islets of Langerhans, Biophys. J. 113 (2017) 1093–1108. https://doi.org/10.1016/j.bpj.2017.07.021.

[20] D. Korošak, M. Jusup, B. Podobnik, A. Stožer, J. Dolenšek, P. Holme, M.S. Rupnik, Autopoietic Influence Hierarchies in Pancreatic β Cells, Phys. Rev. Lett. 127 (2021) 168101. https://doi.org/10.1103/PhysRevLett.127.168101.

[21] P. Chabosseau, G.A. Rutter, S.J. Millership, Importance of Both Imprinted Genes and Functional Heterogeneity in Pancreatic Beta Cells: Is There a Link?, Int. J. Mol. Sci. 22 (2021) 1000. https://doi.org/10.3390/ijms22031000.

[22] R.K.P. Benninger, V. Kravets, The physiological role of β-cell heterogeneity in pancreatic islet function, Nat. Rev. Endocrinol. 18 (2022) 9–22. https://doi.org/10.1038/s41574-021-00568-0.

[23] D. Nasteska, N.H.F. Fine, F.B. Ashford, F. Cuozzo, K. Viloria, G. Smith, A. Dahir, P.W.J. Dawson, Y.-C. Lai, A. Bastidas-Ponce, M. Bakhti, G.A. Rutter, R. Fiancette, R. Nano, L. Piemonti, H. Lickert, Q. Zhou, I. Akerman, D.J. Hodson, PDX1LOW MAFALOW β-cells contribute to islet function and insulin release, Nat. Commun. 12 (2021) 674. https://doi.org/10.1038/s41467-020-20632-z.

[24] V. Kravets, J.M. Dwulet, W.E. Schleicher, D.J. Hodson, A.M. Davis, L. Pyle, R.A. Piscopio, M. Sticco-Ivins, R.K.P. Benninger, Functional architecture of the pancreatic islets reveals first responder cells which drive the first-phase [Ca2+] response, (2021) 2020.12.22.424082. https://doi.org/10.1101/2020.12.22.424082.

[25] D.W. Huang, B.T. Sherman, R.A. Lempicki, Systematic and integrative analysis of large gene lists using DAVID bioinformatics resources, Nat. Protoc. 4 (2009) 44–57. https://doi.org/10.1038/nprot.2008.211.

[26] S.-I. Tschen, S. Dhawan, T. Gurlo, A. Bhushan, Age-dependent decline in beta-cell proliferation restricts the capacity of beta-cell regeneration in mice, Diabetes. 58 (2009) 1312–1320. https://doi.org/10.2337/db08-1651.

[27] E.C. Lewis, C.A. Dinarello, Responses of IL-18-and IL-18 receptor-deficient pancreatic islets with convergence of positive and negative signals for the IL-18 receptor, Proc. Natl. Acad. Sci. U. S. A. 103 (2006) 16852–16857. https://doi.org/10.1073/pnas.0607917103.

[28] M.A. Darwish, A.M. Abo-Youssef, B.A.S. Messiha, A.A. Abo-Saif, M.S. Abdel-Bakky, Resveratrol inhibits macrophage infiltration of pancreatic islets in streptozotocin-induced type 1 diabetic mice via attenuation of the CXCL16/NF-κB p65 signaling pathway, Life Sci. 272 (2021) 119250. https://doi.org/10.1016/j.lfs.2021.119250.

[29] J.F. Garbincius, J.W. Elrod, Mitochondrial calcium exchange in physiology and disease, Physiol. Rev. 102 (2022) 893–992. https://doi.org/10.1152/physrev.00041.2020.

[30] G.A. Rutter, E. Georgiadou, A. Martinez-Sanchez, T.J. Pullen, Metabolic and functional specialisations of the pancreatic beta cell: gene disallowance, mitochondrial metabolism and intercellular connectivity, Diabetologia. 63 (2020) 1990–1998. https://doi.org/10.1007/s00125-020-05205-5.

[31] T.T.-H. Lu, S. Heyne, E. Dror, E. Casas, L. Leonhardt, T. Boenke, C.-H. Yang, Sagar, L. Arrigoni, K. Dalgaard, R. Teperino, L. Enders, M. Selvaraj, M. Ruf, S.J. Raja, H. Xie, U. Boenisch, S.H. Orkin, F.C. Lynn, B.G. Hoffman, D. Grün, T. Vavouri, A.M. Lempradl, J.A. Pospisilik, The Polycomb-Dependent Epigenome Controls β Cell Dysfunction, Dedifferentiation, and Diabetes, Cell Metab. 27 (2018) 1294–1308.e7. https://doi.org/10.1016/j.cmet.2018.04.013.

[32] J. van Arensbergen, J. García-Hurtado, M.A. Maestro, M. Correa-Tapia, G.A. Rutter, M. Vidal, J. Ferrer, Ring1b bookmarks genes in pancreatic embryonic progenitors for repression in adult β cells, Genes Dev. 27 (2013) 52–63. https://doi.org/10.1101/gad.206094.112.

[33] E. Haythorne, M. Rohm, M. van de Bunt, M.F. Brereton, A.I. Tarasov, T.S. Blacker, G. Sachse, M. Silva dos Santos, R. Terron Exposito, S. Davis, O. Baba, R. Fischer, M.R. Duchen, P. Rorsman, J.I. MacRae, F.M. Ashcroft, Diabetes causes marked inhibition of mitochondrial metabolism in pancreatic β-cells, Nat. Commun. 10 (2019) 2474. https://doi.org/10.1038/s41467-019-10189-x.

[34] P. Marchetti, A.M. Schulte, L. Marselli, E. Schoniger, M. Bugliani, W. Kramer, L. Overbergh, S. Ullrich, A.L. Gloyn, M. Ibberson, G. Rutter, P. Froguel, L. Groop, M.I. McCarthy, F. Dotta, R. Scharfmann, C. Magnan, D.L. Eizirik, C. Mathieu, M. Cnop, B. Thorens, M. Solimena, Fostering improved human islet research: a European perspective, Diabetologia. 62 (2019) 1514–1516. https://doi.org/10.1007/s00125-019-4911-4.

[35] R. Arrojo e Drigo, S. Jacob, C.F. García-Prieto, X. Zheng, M. Fukuda, H.T.T. Nhu, O. Stelmashenko, F.L.M. Peçanha, R. Rodriguez-Diaz, E. Bushong, T. Deerinck, S. Phan, Y. Ali, I. Leibiger, M. Chua, T. Boudier, S.-H. Song, M. Graf, G.J. Augustine, M.H. Ellisman, P.-O. Berggren, Structural basis for delta cell paracrine regulation in pancreatic islets, Nat. Commun. 10 (2019) 3700. https://doi.org/10.1038/s41467-019-11517-x.

[36] G.A. Rutter, N. Ninov, V. Salem, D.J. Hodson, Comment on Satin et al. “Take Me To Your Leader”: An Electrophysiological Appraisal of the Role of Hub Cells in Pancreatic Islets. Diabetes 2020;69:830–836, Diabetes. 69 (2020) e10–e11. https://doi.org/10.2337/db20-0501.

[37] A. Stožer, M. Gosak, J. Dolenšek, M. Perc, M. Marhl, M.S. Rupnik, D. Korošak, Functional Connectivity in Islets of Langerhans from Mouse Pancreas Tissue Slices, PLOS Comput. Biol. 9 (2013) e1002923. https://doi.org/10.1371/journal.pcbi.1002923.

[38] E. Akalestou, K. Suba, L. Lopez-Noriega, E. Georgiadou, P. Chabosseau, A. Gallie, A. Wretlind, C. Legido-Quigley, I. Leclerc, V. Salem, G.A. Rutter, Intravital imaging of islet Ca2+ dynamics reveals enhanced β cell connectivity after bariatric surgery in mice, Nat. Commun. 12 (2021) 5165. https://doi.org/10.1038/s41467-021-25423-8.

[39] D.J. Hodson, R.K. Mitchell, E.A. Bellomo, G. Sun, L. Vinet, P. Meda, D. Li, W.-H. Li, M. Bugliani, P. Marchetti, D. Bosco, L. Piemonti, P. Johnson, S.J. Hughes, G.A. Rutter, Lipotoxicity disrupts incretin-regulated human β cell connectivity, J. Clin. Invest. 123 (2013) 4182–4194. https://doi.org/10.1172/JCI68459.

[40] S. Postić, S. Sarikas, J. Pfabe, V. Pohorec, L.K. Bombek, N. Sluga, M.S. Klemen, J. Dolenšek, D. Korošak, A. Stožer, C. Evans-Molina, J.D. Johnson, M.S. Rupnik, High resolution analysis of the cytosolic Ca ^2+^ events in beta cell collectives in situ, Physiology, 2021. https://doi.org/10.1101/2021.04.14.439796.

[41] T.H. Kim, X. Zhou, M. Chen, Demystifying “drop-outs” in single-cell UMI data, Genome Biol. 21 (2020) 196. https://doi.org/10.1186/s13059-020-02096-y.

[42] T. Wang, B. Li, C.E. Nelson, S. Nabavi, Comparative analysis of differential gene expression analysis tools for single-cell RNA sequencing data, BMC Bioinformatics. 20 (2019) 40. https://doi.org/10.1186/s12859-019-2599-6.

[43] W. Hou, Z. Ji, H. Ji, S.C. Hicks, A systematic evaluation of single-cell RNA-sequencing imputation methods, Genome Biol. 21 (2020) 218. https://doi.org/10.1186/s13059-020-02132-x.

[44] J.W. Hughes, J.H. Cho, H.E. Conway, M.R. DiGruccio, X.W. Ng, H.F. Roseman, D. Abreu, F. Urano, D.W. Piston, Primary cilia control glucose homeostasis via islet paracrine interactions, Proc. Natl. Acad. Sci. U. S. A. 117 (2020) 8912–8923. https://doi.org/10.1073/pnas.2001936117.

[45] V. Singla, J.F. Reiter, The Primary Cilium as the Cell’s Antenna: Signaling at a Sensory Organelle, Science. 313 (2006) 629–633. https://doi.org/10.1126/science.1124534.

[46] F. Volta, M.J. Scerbo, A. Seelig, R. Wagner, N. O’Brien, F. Gerst, A. Fritsche, H.-U. Häring, A. Zeigerer, S. Ullrich, J.M. Gerdes, Glucose homeostasis is regulated by pancreatic β-cell cilia via endosomal EphA-processing, Nat. Commun. 10 (2019) 5686. https://doi.org/10.1038/s41467-019-12953-5.

[47] J.H. Cho, Z.A. Li, L. Zhu, B.D. Muegge, H.F. Roseman, E.Y. Lee, T. Utterback, L.G. Woodhams, P.V. Bayly, J.W. Hughes, Islet primary cilia motility controls insulin secretion, Sci. Adv. 8 (2022) eabq8486. https://doi.org/10.1126/sciadv.abq8486.

[48] J.T. Walker, D.C. Saunders, V. Rai, C. Dai, P. Orchard, A.L. Hopkirk, C.V. Reihsmann, Y. Tao, S. Fan, S. Shrestha, A. Varshney, J.J. Wright, Y.D. Pettway, C. Ventresca, S. Agarwala, R. Aramandla, G. Poffenberger, R. Jenkins, N.J. Hart, D.L. Greiner, L.D. Shultz, R. Bottino, Human Pancreas Analysis Program, J. Liu, S.C.J. Parker, A.C. Powers, M. Brissova, RFX6-mediated dysregulation defines human β cell dysfunction in early type 2 diabetes, Cell Biology, 2021. https://doi.org/10.1101/2021.12.16.466282.

[49] J.M. Gerdes, S. Christou-Savina, Y. Xiong, T. Moede, N. Moruzzi, P. Karlsson-Edlund, B. Leibiger, I.B. Leibiger, C.-G. Östenson, P.L. Beales, P.-O. Berggren, Ciliary dysfunction impairs beta-cell insulin secretion and promotes development of type 2 diabetes in rodents, Nat. Commun. 5 (2014) 5308. https://doi.org/10.1038/ncomms6308.

[50] S. Lodh, Primary Cilium, An Unsung Hero in Maintaining Functional β-cell Population, Yale J. Biol. Med. 92 (2019) 471–480.

[51] M. Händel, S. Schulz, A. Stanarius, M. Schreff, M. Erdtmann-Vourliotis, H. Schmidt, G. Wolf, V. Höllt, Selective targeting of somatostatin receptor 3 to neuronal cilia, Neuroscience. 89 (1999) 909–926. https://doi.org/10.1016/S0306-4522(98)00354-6.

[52] T. Iwanaga, T. Miki, H. Takahashi-Iwanaga, Restricted expression of somatostatin receptor 3 to primary cilia in the pancreatic islets and adenohypophysis of mice, Biomed. Res. 32 (2011) 73–81. https://doi.org/10.2220/biomedres.32.73.

[53] A.K. O’Connor, E.B. Malarkey, N.F. Berbari, M.J. Croyle, C.J. Haycraft, P.D. Bell, P. Hohenstein, R.A. Kesterson, B.K. Yoder, An inducible CiliaGFP mouse model for in vivo visualization and analysis of cilia in live tissue, Cilia. 2 (2013) 8. https://doi.org/10.1186/2046-2530-2-8.

[54] F. Ye, D.K. Breslow, E.F. Koslover, A.J. Spakowitz, W.J. Nelson, M.V. Nachury, Single molecule imaging reveals a major role for diffusion in the exploration of ciliary space by signaling receptors, ELife. 2 (2013) e00654. https://doi.org/10.7554/eLife.00654.

[55] J. Luo, Z.-L. Deng, X. Luo, N. Tang, W.-X. Song, J. Chen, K.A. Sharff, H.H. Luu, R.C. Haydon, K.W. Kinzler, B. Vogelstein, T.-C. He, A protocol for rapid generation of recombinant adenoviruses using the AdEasy system, Nat. Protoc. 2 (2007) 1236–1247. https://doi.org/10.1038/nprot.2007.135.

[56] M.A. Ravier, G.A. Rutter, Isolation and Culture of Mouse Pancreatic Islets for Ex Vivo Imaging Studies with Trappable or Recombinant Fluorescent Probes, in: A. Ward, D. Tosh (Eds.), Mouse Cell Cult. Methods Protoc., Humana Press, Totowa, NJ, 2010: pp. 171–184. https://doi.org/10.1007/978-1-59745-019-5_12.

[57] J. Dolenšek, A. Stožer, M. Skelin Klemen, E.W. Miller, M. Slak Rupnik, The Relationship between Membrane Potential and Calcium Dynamics in Glucose-Stimulated Beta Cell Syncytium in Acute Mouse Pancreas Tissue Slices, PLoS ONE. 8 (2013) e82374. https://doi.org/10.1371/journal.pone.0082374.

[58] R. Patro, G. Duggal, M.I. Love, R.A. Irizarry, C. Kingsford, Salmon provides fast and bias-aware quantification of transcript expression, Nat. Methods. 14 (2017) 417–419. https://doi.org/10.1038/nmeth.4197.

[59] M.I. Love, W. Huber, S. Anders, Moderated estimation of fold change and dispersion for RNA-seq data with DESeq2, Genome Biol. 15 (2014) 550. https://doi.org/10.1186/s13059-014-0550-8.

[60] Empirical Mode Decomposition, (n.d.). https://perso.ens-lyon.fr/patrick.flandrin/emd.html (accessed January 19, 2022).

[61] N.E. Huang, S.S.P. Shen, Hilbert–Huang Transform and Its Applications, 2nd ed., WORLD SCIENTIFIC, 2014. https://doi.org/10.1142/8804.

[62] N.E. Huang, INTRODUCTION TO THE HILBERT–HUANG TRANSFORM AND ITS RELATED MATHEMATICAL PROBLEMS, in: Interdiscip. Math. Sci., WORLD SCIENTIFIC, 2005: pp. 1–26. https://doi.org/10.1142/9789812703347_0001.

[63] N.E. Huang, Z. Wu, S.R. Long, K.C. Arnold, X. Chen, K. Blank, On instantaneous frequency, Adv. Adapt. Data Anal. 01 (2009) 177–229. https://doi.org/10.1142/S1793536909000096.

[64] B. Chen, S. Zhao, P. Li, Application of Hilbert-Huang Transform in Structural Health Monitoring: A State-of-the-Art Review, Math. Probl. Eng. 2014 (2014) 1–22. https://doi.org/10.1155/2014/317954.

[65] W.L. Dunn, J.K. Shultis, Exploring Monte Carlo methods, Elsevier, Amsterdam [u.a., 2012.

[66] M. Björnsdotter, K. Rylander, J. Wessberg, A Monte Carlo method for locally multivariate brain mapping, NeuroImage. 56 (2011) 508–516. https://doi.org/10.1016/j.neuroimage.2010.07.044.

[67] wgPlot – Weighted Graph Plot (a better version of gplot) - File Exchange - MATLAB Central, (n.d.). https://www.mathworks.com/matlabcentral/fileexchange/24035-wgplot-weighted-graph-plot-a-better-version-of-gplot (accessed January 19, 2022).

[68] D.J. Hodson, F. Molino, P. Fontanaud, X. Bonnefont, P. Mollard, Investigating and Modelling Pituitary Endocrine Network Function: Modelling Pituitary Network Function, J. Neuroendocrinol. 22 (2010) 1217–1225. https://doi.org/10.1111/j.1365-2826.2010.02052.x.

[69] Power Law, Exponential and Logarithmic Fit - File Exchange - MATLAB Central, (n.d.). https://www.mathworks.com/matlabcentral/fileexchange/29545-power-law-exponential-and-logarithmic-fit (accessed January 19, 2022).

[70] GitHub - michellehirsch/xlswritefig: Write a MATLAB figure to an Excel spreadsheet, (n.d.). https://github.com/michellehirsch/xlswritefig (accessed January 19, 2022).

[71] K. Sameshima, L.A. Baccala, eds., Methods in Brain Connectivity Inference through Multivariate Time Series Analysis, CRC Press, Boca Raton, 2014. https://doi.org/10.1201/b16550.

[72] E. Zivot, J. Wang, Vector Autoregressive Models for Multivariate Time Series, in: Model. Financ. Time Ser.-Plus^®^, Springer New York, New York, NY, 2003: pp. 369–413. https://doi.org/10.1007/978-0-387-21763-5_11.

[73] C.W. Granger, Investigating causal relations by econometric models and cross-spectral methods, Econom. J. Econom. Soc. (1969) 424–438.

[74] E.E. Leamer, Vector autoregressions for causal inference?, Carnegie-Rochester Conf. Ser. Public Policy. 22 (1985) 255–304. https://doi.org/10.1016/0167-2231(85)90035-1.

[75] K.J. Blinowska, M. Kamiński, Multivariate Signal Analysis by Parametric Models, in: Handb. Time Ser. Anal., John Wiley & Sons, Ltd, 2006: pp. 373–409. https://doi.org/10.1002/9783527609970.ch15.

[76] Vector Autoregression (VAR) - Comprehensive Guide with Examples in Python, Mach. Learn. Plus. (2019). https://www.machinelearningplus.com/time-series/vector-autoregression-examples-python/ (accessed May 11, 2022).

[77] Econometrics Beat: Dave Giles’ Blog: Testing for Granger Causality, (n.d.). https://davegiles.blogspot.com/2011/04/testing-for-granger-causality.html (accessed May 11, 2022).

[78] P. Pedroni, CRITICAL VALUES FOR COINTEGRATION TESTS IN HETEROGENEOUS PANELS WITH MULTIPLE REGRESSORS, (1999) 18.

[79] R. Mushtaq, Augmented Dickey Fuller Test, Social Science Research Network, Rochester, NY, 2011. https://doi.org/10.2139/ssrn.1911068.

[80] M.W. Wright, T.A. Eyre, M.J. Lush, S. Povey, E.A. Bruford, HCOP: the HGNC comparison of orthology predictions search tool, Mamm. Genome Off. J. Int. Mamm. Genome Soc. 16 (2005) 827–828. https://doi.org/10.1007/s00335-005-0103-2.

[81] T.A. Eyre, M.W. Wright, M.J. Lush, E.A. Bruford, HCOP: a searchable database of human orthology predictions, Brief. Bioinform. 8 (2007) 2–5. https://doi.org/10.1093/bib/bbl030.

[82] R.L. Seal, S.M. Gordon, M.J. Lush, M.W. Wright, E.A. Bruford, genenames.org: the HGNC resources in 2011, Nucleic Acids Res. 39 (2011) D514–519. https://doi.org/10.1093/nar/gkq892.

[83] J. Tabak, Geometry: The Language of Space and Form, Facts On File, Incorporated, 2014. https://books.google.co.uk/books?id=r0HuPiexnYwC.

[84] J.N. Hansen, S. Rassmann, B. Stüven, N. Jurisch-Yaksi, D. Wachten, CiliaQ: a simple, open-source software for automated quantification of ciliary morphology and fluorescence in 2D, 3D, and 4D images, Eur. Phys. J. E. 44 (2021) 18. https://doi.org/10.1140/epje/s10189-021-00031-y.

