## Supplementary figures and images for "Molecular phenotyping of single pancreatic islet leader beta cells by “Flash-Seq”"

### Supplemental Figures

#### Supplementary Figure 1

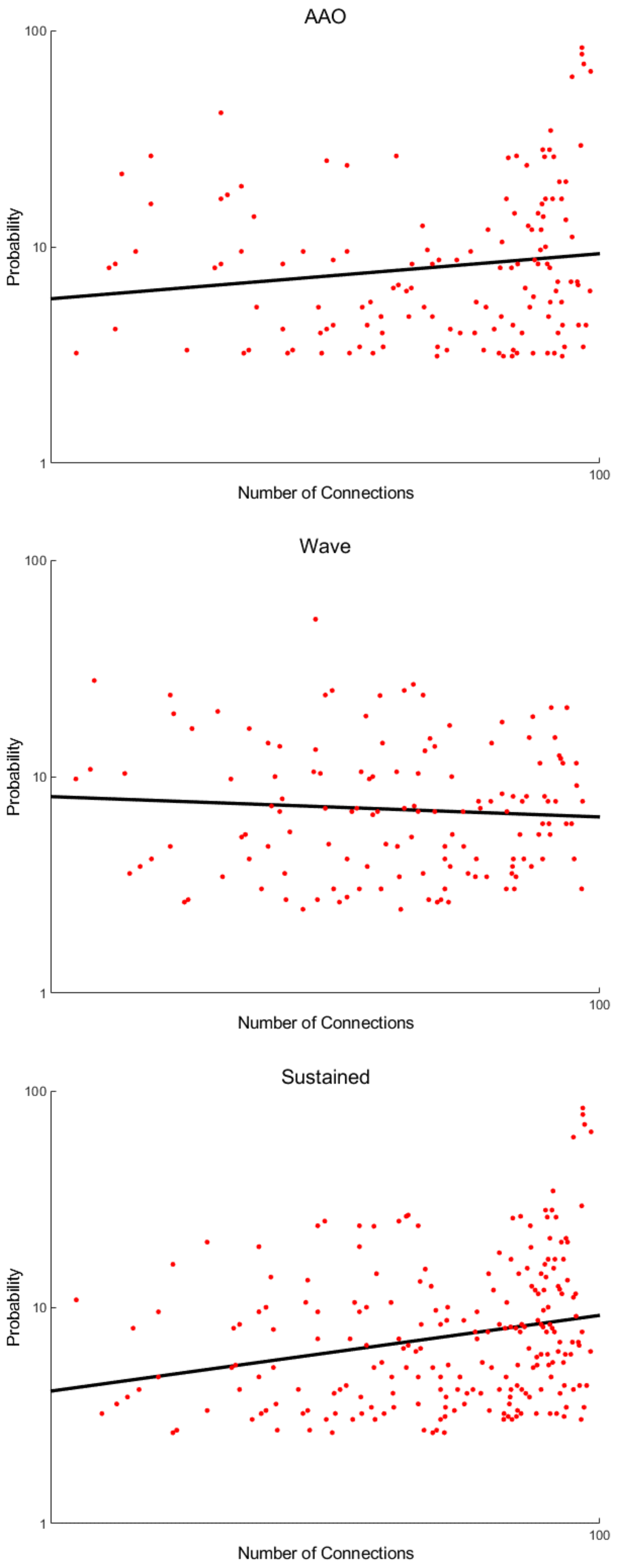

Supplementary Figure 2

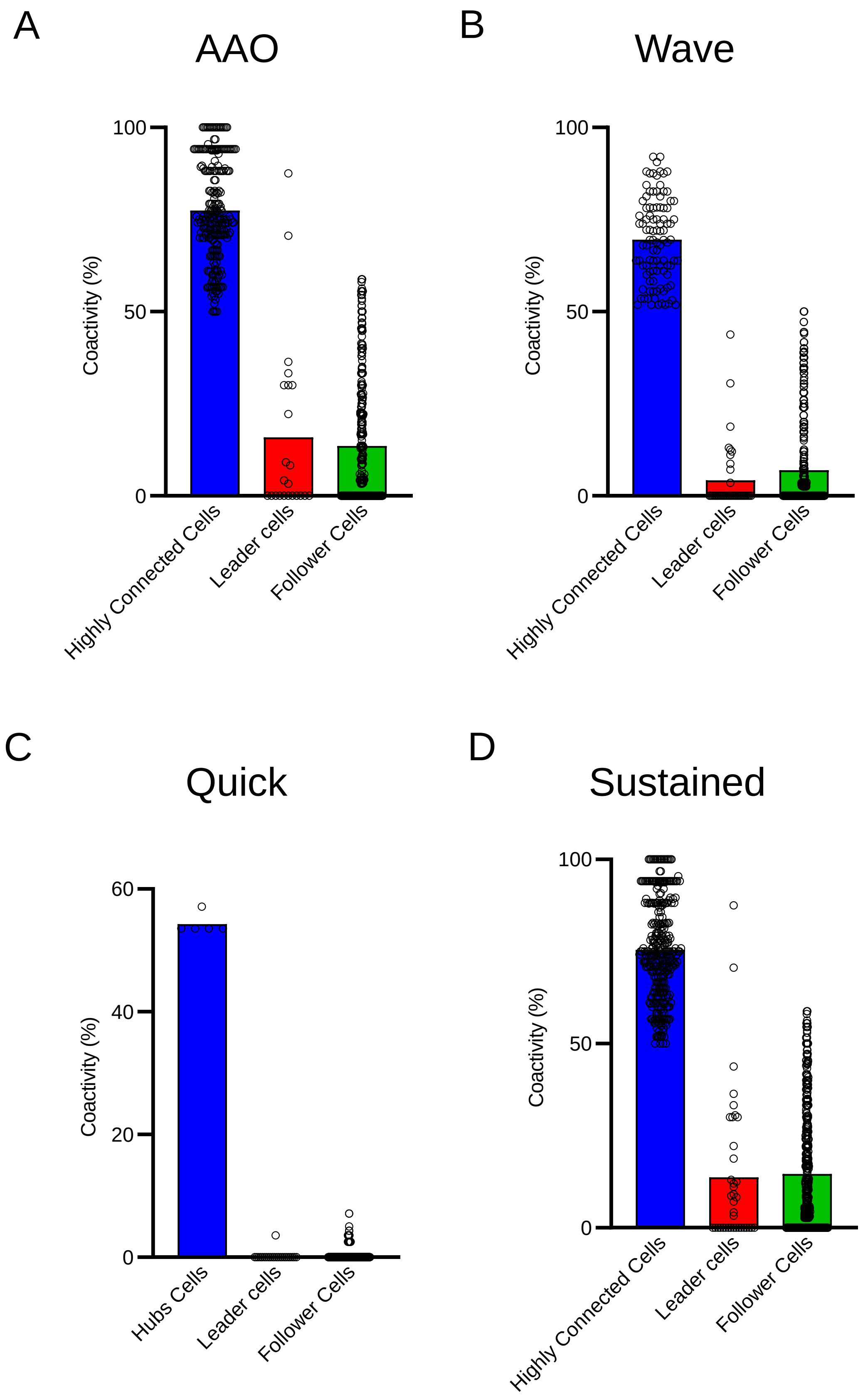
